# Zona Incerta activity dynamics underlying associative fear learning and fear generalization

**DOI:** 10.1101/2021.07.31.454576

**Authors:** Zhuoliang Li, Giorgio Rizzi, Kelly R. Tan

## Abstract

Recent studies suggest that the Zona Incerta (ZI) plays a role in fear learning and recall. However, there is a clear gap in knowledge as to whether the ZI can encode fear evoking threats and cues that predict them. Here, we subject mice to a classical fear conditioning paradigm while recording the *in-vivo* calcium dynamics of ZI neurons. We observed that ZI neurons can encode not only a fear evoking stimulus, but can also learn to encode predictive cues, associate them with the unconditioned fear evoking stimulus, and discriminate them from neutral non-predictive cues. While the ZI across all mice learned to become mainly excited by fear predictive cues, only mice that did not generalize fear became largely inhibited by non-predictive cues and better discriminated the two. Together, we provide extensive evidence that the ZI differentially encode fear predictive and non-predictive cues in a generalization dependent manner.

## Introduction

Fear is an emotional defensive response to a threatening stimulus^1, 2^. To ensure survival, it is important to react not only to innately threatening stimuli but also those that predict them^1–4^. A wide range of brain regions are involved in this learned fear response^3^ including the amygdala^5, 6^, prefrontal cortex^7, 8^, hippocampus^9, 10^, lateral ventral tegmental area^11^, and periaqueductal gray^12^. Recently, the ZI has also be implicated in fear. Specifically, optogenetic inactivation of the GAD2-expressing ZI neurons in mice^13^ and deep brain stimulation of the ZI in patients with Parkinson’s Disease^14^ improved fear recognition. Alternatively, the irreversible tetanus toxin mediated blockade of synaptic transmission of all ZI neurons or only parvalbumin expressing ones impaired fear learning and recall^15^. Generalization is an aspect of learned fear where neutral and similar cues are also considered threatening and evoke the learned fear response^16^. Interestingly, the chemogenetic activation and inactivation of ZI neurons has been shown to potentiate and reduce fear generalization during recall, respectively^17^. In spite of the contentious roles of the ZI in fear and fear generalization, little is known about whether ZI neurons can encode fear associated cues and fear generalization.

## Results

### Zona Incerta cells differentially encode fear predictive and non-predictive cues

To investigate whether the ZI neurons can encode fear associated cues, we recorded the calcium dynamics of ZI neurons as a proxy for neuronal activity while mice performed in a classical fear conditioning paradigm. We surgically injected a virus containing the calcium indicator GCaMP6m (AAV1-CAMKIIα-GCaMP6m) into the ZI and implanted a gradient refractive index (GRIN) lens above (Fig. 1A). The calcium transients of ZI neurons were concomitantly imaged while mice were conditioned to a pre-exposed auditory tone (CS+) followed with FS and another without (CS−) (Fig. 1B). Mice exhibited similarly low levels of freezing to the neutral context as well as the conditioning context before any shocks were delivered and learned to freeze to the conditioning context (Fig. 1C). CS+ evoked low and similar levels of freezing to the CS− before conditioning and significantly more freezing on the test day (Video S1-3, Fig. 1D), hence showing that they have learned that the CS+ is predictive of the FS. Interestingly, cue related increases and decreases in calcium transients were observed in ZI neurons throughout the task (Video S4, Fig. 1A). To characterize these responses, we compared the activity during the cue window against the same duration (baseline) before (ranksum test, p< 0.05). Additionally, we calculated the discrimination index (DI) assessing the likelihood that activity during the cue window can be discriminated from the baseline with the receiver-operating characteristic analysis. To investigate whether these cue related responses can be distinguished from spontaneous changes in activity, we compared each DI with DIs calculated from randomly sampled time windows (bootstrap test, α = 0.05). While ZI neurons bi-directionally responded to both the CS− and CS+ throughout the task, the majority of responsive cells were inhibited by the CS− (Fig. 1E-H) and excited by the CS+ (Fig. 1I-L) after learning. Proportionally, more responses were distinct from the spontaneous activity after learning (Fig 1H,L). Furthermore, the amplitude and discriminability of the inhibition to the CS− (Fig. 1M-N) and excitation to the CS+ (Fig. 1O-P) became increasingly potentiated throughout learning and peaked on the test day. Together, these findings suggest that the ZI learns to become excited and inhibited by fear predictive and non-predictive cues, respectively.

**Figure 1.**
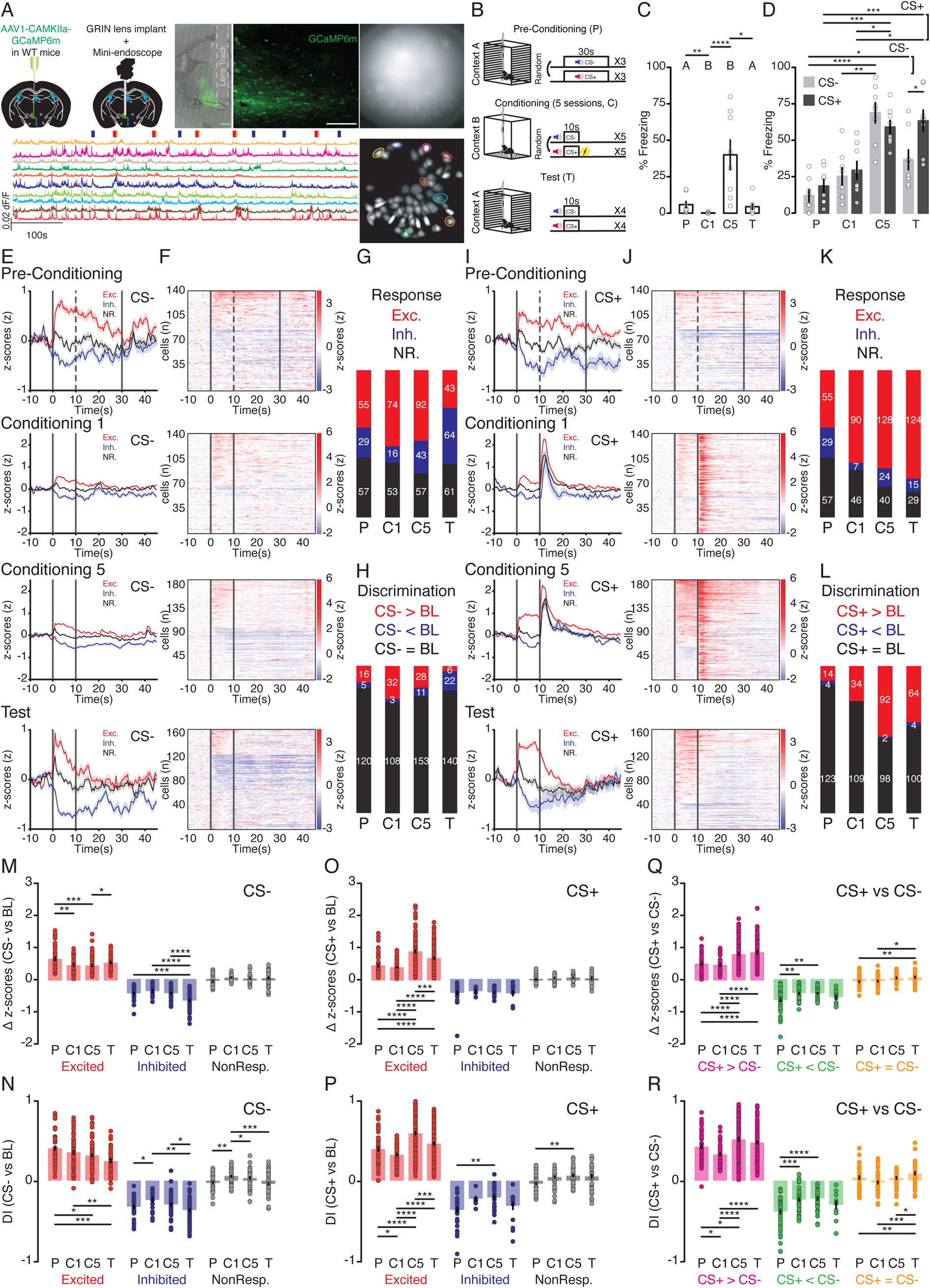
ZI neurons are bi-directionally modulated by auditory cues and are plastic across over fear conditioning. A) Design to visualize calcium dynamics in ZI neurons: injection of GCaMP6m containing virus, GRIN lens implant, low and high magnification confocal image of ZI containing slice with GCaMP6m expression in the ZI and lens track above (500 and 100µm scale bar, respectively), raw and maximal projection images of field-of-view with example traces. B) Fear conditioning paradigm. C) Bar graph of contextual fear across P, C1, C5, and T. D) Bar graph of cued fear across P, C1, C5, and T. E) Average traces (± SEM, shaded) of categorized cells responding to the CS− (black bars) across P, C1, C5, and T. F) Heatmap of CS− responses across P, C1, C5, and T. G) Stacked bar chart depicting the proportion of categorized responses to CS− across P, C1, C5, and T. H) Stacked bar chart showing the proportion of CS− responses that can be discriminated from intrinsic fluctuations within the whole trace across P, C1, C5, and T. I) Average traces of categorized cells responding to the CS+ across P, C1, C5, and T. J) Heatmap of CS+ responses across P, C1, C5, and T. K) Stacked bar chart depicting the proportion of categorized responses to CS+ across P, C1, C5, and T. L) Stacked bar chart with the proportion of CS+ responses that can be discriminated from intrinsic fluctuations across P, C1, C5, and T. M) Bar graph of the amplitude of CS− responses across P, C1, C5, and T. N) Bar graph of the DI of CS− responses across P, C1, C5, and T. O) Bar graph of the amplitude of CS+ responses across P, C1, C5, and T. P) Bar graph of the DI of CS+ responses across P, C1, C5, and T. Q) Bar graph of the amplitude difference between CS+ and CS− responses across P, C1, C5, and T. R) Bar graph of the DI between CS+ and CS− responses across P, C1, C5, and T. Data are presented as mean ± SEM *p<0.05, **p<0.01, ***p<0.001, ****p<0.0001, 141-192 cells from 8 mice.

The sharp contrast in the way the majority of ZI neurons responded to the CS− and the CS+ after learning led us to speculate that they discriminate between the two tones. To test this hypothesis, we compared the activity of ZI neurons between the two tones (Fig. S1A-D). While many ZI neurons responded differently to the CS− and the CS+ even before conditioning (Fig. S1C), the majority of these differences were indistinguishable from spontaneous changes in activity (Fig. S1D). After learning, the majority of cells exhibited higher activity during the CS+ than the CS− (Fig. S1C) and a noticeable portion of these differences now distinguishable from spontaneous activity changes (Fig. S1D). In addition, this shift coincided with an increase in the differences in amplitude (Fig. 1Q) and discriminability (Fig. 1R) between the CS+ and the CS− windows within these cells.

### Zona Incerta cells are mostly excited by footshocks

In addition to predictive cues, we also investigated whether ZI neurons are modulated by the FS (Fig. S1E-M). We observed that the majority of ZI neurons were strongly excited by the FS (Fig. S1G) and can discriminate them from spontaneous activity changes (Fig. S1H) on both the first and last conditioning days. Lastly, to examine whether ZI neurons can associate predictive cues with FS, we investigated the similarity between CS+ and FS responses by calculating the Euclidean distance (ED) between the two^11, 18^. Indeed, the ED between CS+ and FS responses became smaller (thus more similar) across pairings (Fig. S1K-L) on both conditioning days, suggesting an association between the two responses. However, the lack of any change in the ED between CS+ and FS+ responses across the first and last conditioning days (Fig. S1M) suggests that this association may not be consolidated. So far, we have observed on a population level across all recorded cells that ZI neurons can encode predictive cues as well as FS, discriminate between CS+ from CS−, and associate CS+ with FS.

### Zona Incerta cells become excited by fear predictive cues by consolidating their association to footshocks

To strengthen our findings, we identified longitudinally tracked cells with spatial alignment (Fig. 2A) and investigated changes in their response throughout fear learning. We classified cells based on their responses to cues on the last day they were presented (T for CS−/CS+ and C5 for FS) and examined these responses previously. Expectedly, we observed similar proportion of longitudinally tracked cells responsive to CS− (Fig. 2B-E) and CS+ (Fig. 2F-I) on the test day as compared to the populational analyses (Fig. 1). Notably, there was a great deal of heterogeneity and plasticity in how cells changed their responses to the CS− and CS+ across days (Fig 2D-E, H-I). Interestingly, the amplitude (Fig. 2L) and DI (Fig. 2M) of the average excitatory responses to CS+ increased gradually across fear learning and peaked during the test day, coinciding with a gradual increase in freezing to the CS+ (Fig. 1D). On the other hand, the corresponding changes to the average inhibitory responses to the CS− were more drastic and appeared only on the test day (Fig. 2B, J-K).

**Figure 2.**
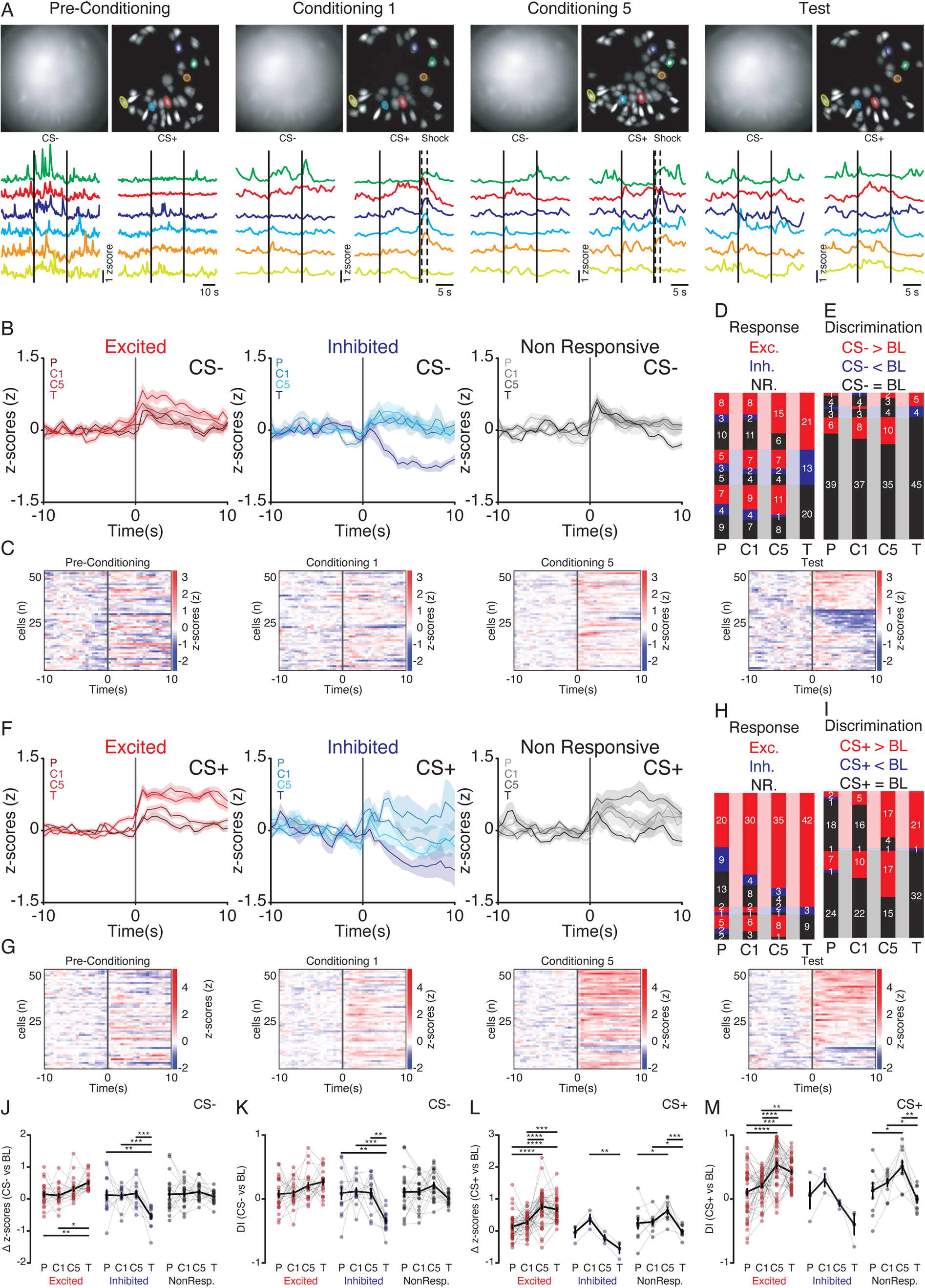
Fear learning gradually potentiate ZI excitatory responses to predictive cues. A) Example longitudinally tracked cells across P, C1, C5, and T: example raw and maximal projection images with example cell traces (black bars represent tones and shock windows). B) Average traces of T day CS− responsive cells tracked across P, C1, C5, and T. C) Heatmaps of tracked CS− responses across P, C1, C5, and T. D) Stacked bar chart of the proportion of CS− responsive cells on the T day and their response profiles before. E) Stacked bar chart with the proportion of CS− responses that can be discriminated from spontaneous activity across P, C1, C5, and T. F) Average traces of T day CS+ responsive cells tracked across P, C1, C5, and T (black bar denotes tone onset). G) Heatmaps of tracked CS+ responses across P, C1, C5, and T. H) Stacked bar chart showing the proportion of CS+ responsive cells on the T day and how they responded days prior. I) Stacked bar chart depicting the proportion of CS+ responses that can be discriminated from intrinsic cell activity across P, C1, C5, and T days. J) Line graph summarizing the tracked changes in amplitude of CS− responses across P, C1, C5, and T days. K) Line graph showing the tracked changes in the DI of CS− responses across P, C1, C5, and T days. L) Line graph depicting the tracked changes in the amplitude of CS+ responses across P, C1, C5, and T days. M) Line graph with the tracked DI of CS+ responses across P, C1, C5, and T days. Data are presented as mean ± SEM *p<0.05, **p<0.01, ***p<0.001, ****p<0.0001, 54 cells from 5 mice.

We also investigated how ZI cells changed their discrimination between the CS+ and CS− across learning. On average, cells responded similarly to the CS+ and CS− (Fig. 3A-F) across fear conditioning and only clearly distinguishable from one another on the test day (Fig. 3E-F), coinciding with the differences in freezing between the CS+ and CS− across the paradigm (Fig. 1D). Furthermore, we also observed in longitudinally tracked cells that the majority of ZI neurons remained stably excited to FS (Fig. 3G-O) and these responses became more similar to the CS+ within conditioning days (Fig. 3M-N) but not between the first and last conditioning day (Fig. 3O). Strikingly, the CS+ responses only became similar to the FS between the first and last conditioning days in cells that learned to respond to the CS+ on the test day (Figure 3P-Q), suggesting that ZI cells differentially consolidate CS+ with FS. Together, these data in both the populational level and in longitudinally tracked cells provide convincing evidence that the ZI neurons encode threatening stimuli, can learn to encode cues that predict them, and discriminate fear-predictive cues from neutral ones.

**Figure 3.**
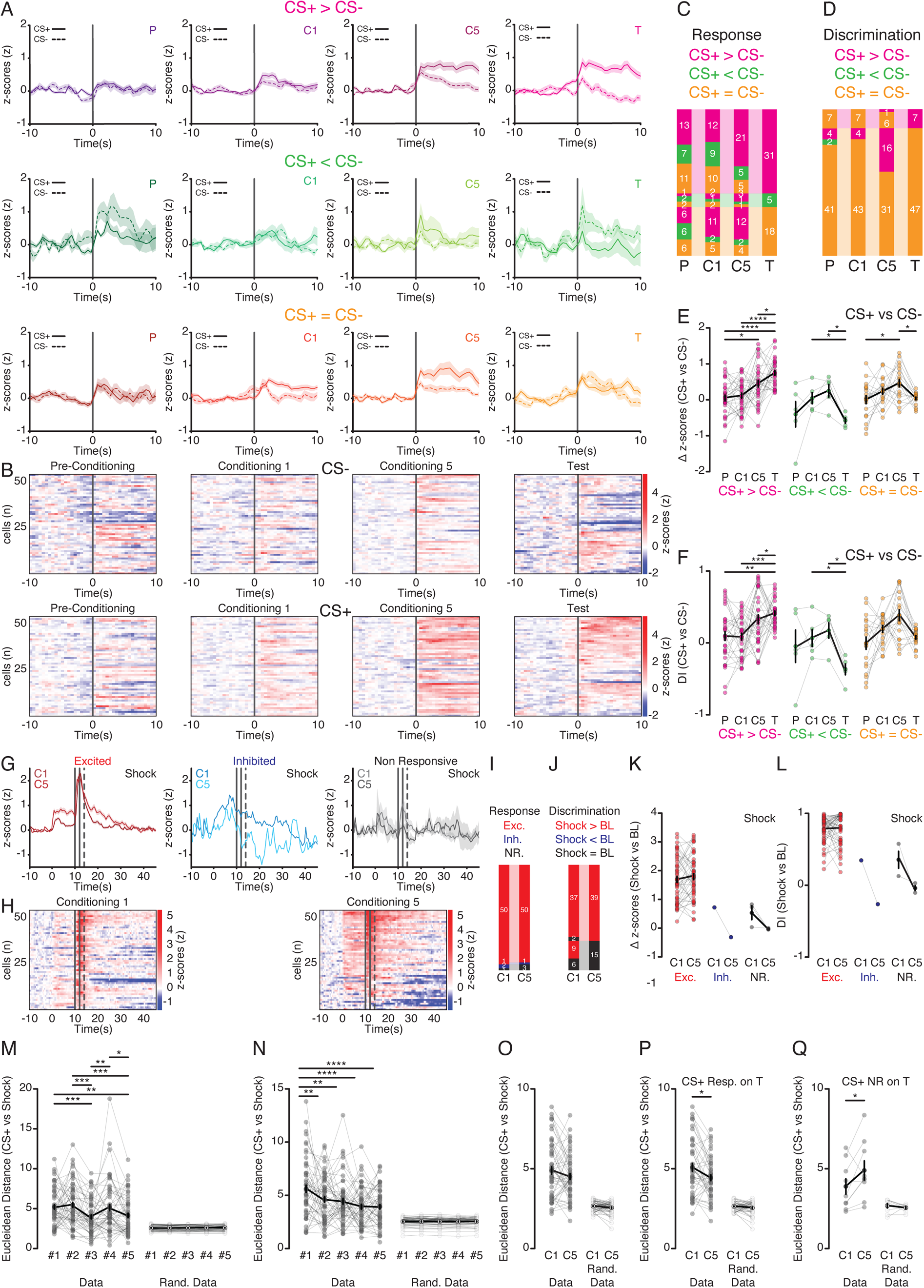
ZI neurons associate CS+ with footshocks and learn to discriminate between CS+ from CS−. A) Average traces of cells that are differentially modulated by CS+ and CS− on the T day tracked across P, C1, C5, and T days (black bar denote tone onset). B) Heatmaps of tracked responses to CS+ and CS− across P, C1, C5, and T. C) Stacked bar chart of proportion of cells that are differentially modulated by CS+ and CS− during the T day and their bias during the days before. D) Stacked bar chart showing the proportion of cells whose discrimination between CS+ and CS− is grossly different from disparity among spontaneous activity across P, C1, C5, and T days. E) Line graph summarizing the tracked amplitude difference between CS+ and CS− responses across P, C1, C5, and T days. F) Line graph depicting the tracked changes in DI between CS+ and CS− across P, C1, C5, and T days. G) Average traces of shock responsive cells on the C5 and their shock responses also during the first conditioning day (solid and dotted bar denotes shock and shock analysis windows, respectively). H) Heatmaps of tracked shock responses across C1 and C5. I) Stacked bar chart with the proportion of shock responsive cells on C5 and their response profile on C1. J) Stacked bar chart of the proportion of shock responses that can be discriminated from intrinsic trace fluctuations across C1 and C5. K) Line graph summarizing the tracked changes in the amplitude of shock responses across C1 and C5. L) Line graph with the tracked changes in the DI of shock responses across C1 and C5. M) Line graph with the Euclidean Distance between the CS+ and FS pairings during C1. N) Line graph of the Euclidean Distance between the CS+ and FS pairings on C5. O) Line graph depicting the average Euclidean Distance between the CS+ and FS pairings tracked across C1 and C5. P) Line graph depicting the average Euclidean Distance between the CS+ and FS pairings in cells that responded to the CS+ on T tracked across C1 and C5. Q) Line graph depicting the average Euclidean Distance between the CS+ and FS pairings in cells that were non-responsive to the CS+ on T tracked across C1 and C5. Data are presented as mean ± SEM *p<0.05, **p<0.01, ***p<0.001, ****p<0.0001, 54 cells from 5 mice.

### Fear generalization impairs ZI’s ability to discriminate between fear predictive and non-predictive cues

Individual differences in fear acquisition and recall have been well documented in humans^19^ and to a lesser extent in rodents^20–22^. It is still unclear whether the activity of ZI neurons is influenced by the extent of fear learning that took place. Given that the ZI has been reported to modulate fear generalization^17^, we therefore investigated whether the fear generalization impacted the learned ZI responses to fear predictive and non-predictive cues. To do so, we stratified our data (Fig. 4, S2-4) by the extent the imaged mice generalized fear based on the differences in freezing between the CS+ and CS− on the test day (fear generalized FG mice, < 20% delta CS+ to CS− freezing, fear not generalized FnG mice, > 20%). Despite similar contextual freezing levels throughout (Fig. 4A), FnG mice better distinguished the CS+ from the CS− after learning (VideoS5-6, Fig. 4B), reflective of lower modality-specific fear generalization. While ZI neurons in both groups were mostly excited by FS (Fig. S2) and by the CS+ after fear learning, ZI neurons only in the FnG mice were mainly inhibited by the CS− (Fig S3). Consequently, proportionally more ZI neurons from FnG mice showed higher activation by the CS+ than the CS− after fear learning in a manner that is distinguishable from the spontaneous activity changes (Fig. 4E-G, S4). Overall, these data demonstrates that fear generalization impairs the way ZI neurons learned to discriminate between fear predictive and non-predictive cues.

**Figure 4.**
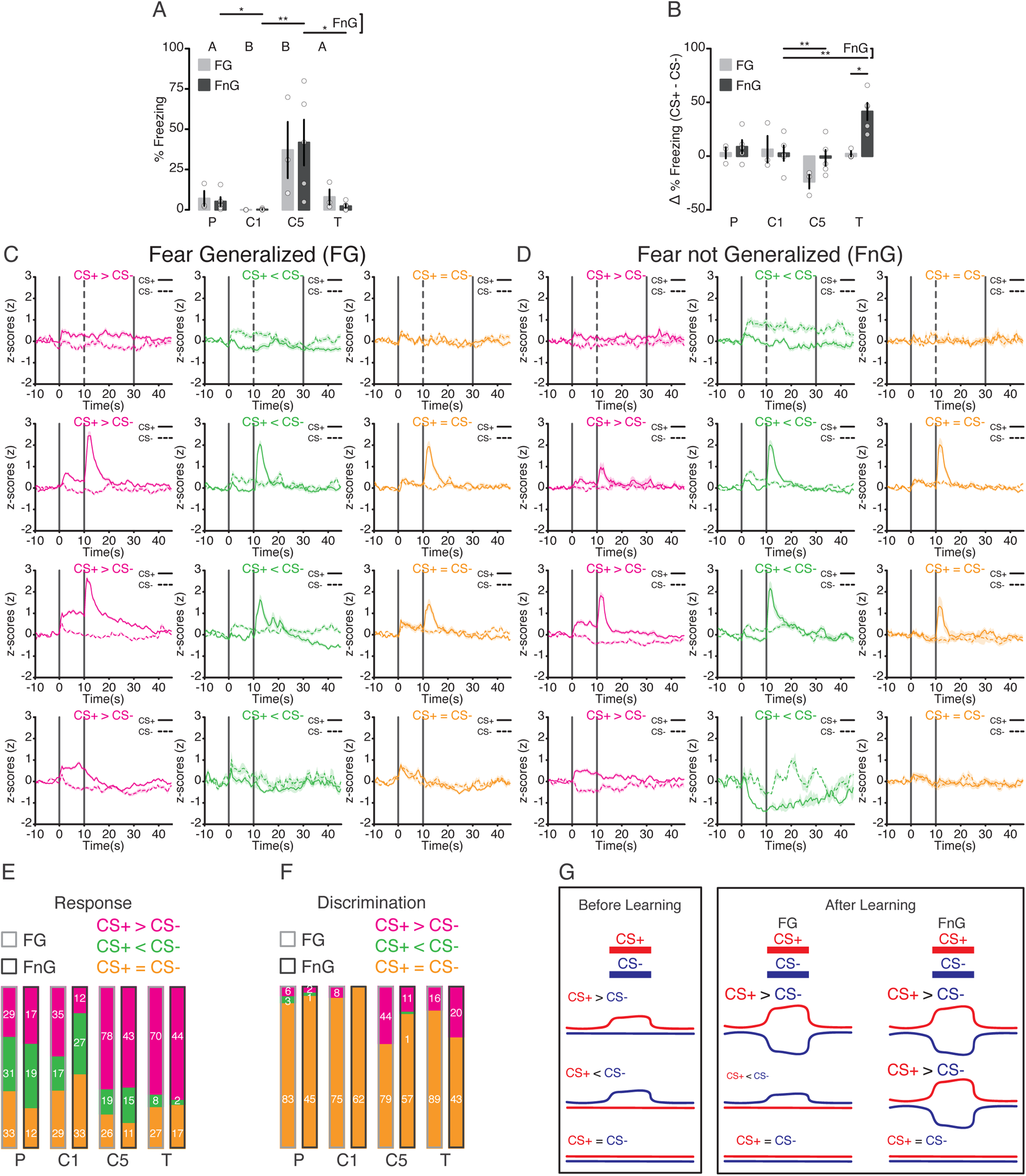
ZI neurons in fear generalized and not generalized mice differentially learn to discriminate between CS+ and CS−. A) Bar graph of contextual fear between fear generalized and not generalized mice across P, C1, C5, and T. B) Bar graph with delta (CS+ minus CS−) tone freezing in fear generalized and not generalized mice across P, C1, C5, and T. C) Average traces of cells differentially modulated by CS+ and CS− in fear generalized mice across P, C1, C5, and T. D) Average traces of cells that respond differentially to CS+ and CS− in mice that did not develop fear generalization across P, C1, C5, and T. E) Stacked bar chart showing the proportion of cells differentially modulated by CS+ and CS− in fear generalized and not generalized mice across P, C1, C5, and T. F) Stacked bar chart depicting the proportion of cells whose discrimination between CS+ and CS− are significantly different from that of intrinsic activity fluctuations in fear generalized and not generalized mice across P, C1, C5, and T. G) Model highlighting the differences in CS+ and CS− discrimination in fear generalized and not generalized mice following fear conditioning. Data are presented as mean ± SEM ^p = 0.0572 (trend), *p<0.05, **p<0.01, ***p<0.001, ****p<0.0001, 81-123 cells (fear generalized, 3 mice) and 48-69 cells (fear not generalized, 5 mice).

## Discussion

Despite a growing number of recent studies implicating a role for the ZI in associative fear learning^13–15, 17, 23^, little is known about how fear learning and the extent of fear generalization impact the way ZI encodes fear predictive and non-predictive cues. By imaging ZI neurons while mice underwent a classical fear conditioning task, we observed that the majority of the ZI neurons are stably excited by the threatening and fear evoking FS, supporting claims that the ZI is sensitive to stressful stimuli^24^. ZI cells associate the CS+ with the FS and become gradually more excited after learning, coinciding with a gradual increase in CS+ evoked freezing. This finding is in conflict with a previous report where no plasticity was observed in the *in vivo* extracellular activity of ZI cells to successive presentations of CS+ during fear conditioning^13^. However, the previous study only reported the absolute activity observed across CS+ windows without taking into account the activity patterns immediately before and how CS+ modulated them. Furthermore, the low number of units recorded (only 12 total) and differences in the behavioral task (ie. having the mice restrained in a head-fixed position, only one conditioning session) could also contribute to the discrepancy. In sharp contrast to the CS+, ZI cells learn to become inhibited by the CS− and consequently, mainly discriminate between the CS+ and the CS− by exhibiting higher activity towards the CS+.

Our findings closely support studies where loss-of-function manipulations in the ZI impaired freezing to fear predictive cues during recall^15, 23^ rather than the report where the inactivation of ZI GAD2-expressing cells enhanced freezing^13^. Future studies should focus on the gain-of-function side and investigate whether entraining the activity of the ZI to cues in the absence of aversive stimuli is sufficient to evoke cued fear.

Overall, our observations suggest that innate and perceived (fear-predictive) threats activate ZI neurons coinciding with an elevated defensive freezing response while neutral cues suppress ZI neuronal activity. Speculatively, a general increase in the excitability of ZI neurons would reduce the effect size of CS+ excitation and potentiate CS− inhibition, consequently resulting in a reduced threat perception and thereby a reduction in freezing to both CS+ and CS−. Conversely, a general decrease in the excitability would enhance threat perception and freezing to both cues. Indeed, these scenarios were observed when the ZI was chemogenetically manipulated prior to fear recall^17^, supporting our theory. Based on our theory, fear generalization would enhance the threat perception and ZI responses to neutral cues. Indeed, we observed more ZI neurons were excited by the CS− in the FG mice than the FnG ones and showed less robust discrimination between the CS+ and CS− both in terms of activity and defensive freezing. Excess fear generalization is associated with post-traumatic stress disorders and generalized anxiety^25, 26^. Future studies should investigate whether the ZI is excessively activated by triggers perceived to be threatening in these pathological conditions and whether the ZI would be an effective therapeutic target.

## Methods

### Animals

Adult (6-8 weeks old) male C57BL/6 mice were bred in house and kept in a temperature-controlled facility under a 12-hour light/dark cycle (7am to 7pm, light period) with standard chow and water provided ad libitum. All experimental procedures were conducted during the light period. All experimental procedures were approved by the Institutional Animal Care Office of the University of Basel and the Cantonal Veterinary Office of Basel under the license number 2742.

### Surgery

Mice were anesthetized with Isoflurane (5% for induction, 1.5% for maintenance, Provet Healthcare, EZ Anesthesia Systems) in O_2_ and placed onto the stereotaxic frame (World Precision Instruments). The eyes were covered by a custom mixed eye lube composed of vegetable oil and petroleum jelly (Vaseline). Lidocaine (0.2mg/kg, Stueli Pharma) was injected (subcutaneous) above the skull. The skin was disinfected through successive applications of Betadine (Mundipharma) and ethanol (70%, Sigma-Aldrich). Hair above the skull were shortened with surgical scissors (Fine Science Tools). Incisions were made to expose the skull and then thoroughly cleaned with hydrogen peroxide (3%, Sigma-Aldrich), ethanol, and saline (Braun). The skull was etched with surgical blades (Swann-Morton) before drilling holes above the ZI (1.3mm posteriorly to the Bregma, 0.7mm laterally to the midline, 4.9mm below the surface of the skull^27^). Viral construct containing the GCaMP6m (AAV1-CaMKIIα-GCaMP6m-WPE-SV40, Ready to Image Virus, Inscopix) was injected (250nl) into the ZI with a piston operated injector system (Narishige). Mice were stitched with tissue absorptive silk sutures (SABANNA) and allowed to recover for 2 weeks before implantation. The mice were prepared as previously described. Two small holes were drill anterior to the Bregma and screws were inserted. A needle (26-guage, Braun) was stereotaxically lowered (0.1mm) above the ZI and then a GRIN (0.6mm diameter, 7.3mm long, Inscopix) lens already integrated with a baseplate was implanted at the same location. A custom metal head-bar (0.4cm wide, 2cm long) was placed on the side and pointing to the posterior direction. The lens was fixed to the skull with a combination of adhesive (Pattex) and light-curable (Kulzer, Kerr) glue and secured together with the head-bar to the skull with a fixed headcap built from dental cement (Kulzer). Buprenorphine (Reckitt Benckiser) was administered (intraperitoneal, 0.1g/kg) as necessary to alleviate pain.

### Fear Conditioning with Calcium Imaging

Implanted were checked bi-weekly until the area below the lens have cleared and GCaMP6m labeled cells can be visualized. To do so, mice were held by the head-bar on a custom-built running wheel while the miniscope (Inscopix) is attached. ZI calcium transients were recorded by the nVoke2 and nVista2 (Inscopix) systems and visualized through the Inscopix Data Acquisition Software (IDAS). Once cells could be visualized, mice were individually handled and habituated to the experimental room (5 mins per day for 3 consecutive days). Following the last handling day, mice were subjected to a 7-day fear conditioning protocol (pre-conditioning, 5 conditioning days, and test day). On each day, the miniscope is first attached and mice were then placed into either a plexiglass box (20cm wide, 25cm long, 15cm tall) covered with polka-dots (context A, pre-conditioning/test days, cleaned with 1% acetic acid) or a different (context B, conditioning days, cleaned with 70% ethanol) plexiglass box (20cm wide, 22.5cm long, 11.5cm tall) with metal gridded floors (Med Associates Inc) and bedding below (replaced between each animal). In each session, a baseline period was first recorded to measure the contextual freezing. Auditory tones (7kHz CS−, 12kHz CS+) were presented (85 dB all tones, 30s/3 times each in pre-conditioning day and 10s/5 times each for conditioning days and 10s/4 times each on the test day) in a randomized order (except for the test day where CS− are presented consecutively and followed by CS+) with a randomized inter-tone interval (ITI) with a 100s average duration (70-130s for pre-conditioning and test days and 50-150s for conditioning days). The duration, number of tones, and ITI were designed to vary and randomized when feasible to reduce expectation and the likelihood that mice attribute either of these factors as fear predictive. Shocks (2s, 0.6mA) were delivered immediately following the CS+ presentations during the conditioning days. These cues were controlled and delivered by ANYMAZE (Stoeling). Videos were captured by a camera (The Imaging Source) placed from above (or at a 45-degree angle) at a fixed sampling rate (30Hz) and freezing was detected with ANYMAZE (1s minimum). Calcium transients were recorded by IDAS. The synchronization between the behavioral and calcium recordings were made possible by the delivery of a short TTL (time-to-live) pulse at the start and end of the fear conditioning protocol and is recorded by the IDAS. Timestamps for all CS− and CS+ presentations are recorded in ANYMAZE and also by IDAS through TTL pulses delivered by ANYMAZE. The resulting files with the timestamped freezing detection, as well as the behavioral and calcium recordings were exported for further analyses.

### Histology

Mice were anesthetized with a lethal injection (intraperitoneal) of pentobarbital (0.3mg/kg, Stueli Pharrma). Mice were laid flat on a styrofoam board and pinned down with needles (32G, Braun). Incisions were made to expose the heart. Mice were then perfused with a cold phosphate buffered saline (PBS, Sigma, 25mL) solution and equal volume of 4% paraformaldehyde (PFA, Sigma) fixative in PBS. The head was removed and post-fixed for at least 48 hours in the PFA fixative. The brain was then carefully extracted and immersed in 30% sucrose (Sigma) in PBS until complete submersion. Sections were prepared with a cryostat (Leica 1950CM) and mounted on glass slides (Superfrost Plus, Thermo Scientific) in a DAPI containing mounting solution (ProLong God Antifade, Invitrrogen) and covered with borosilicate cover glass (VWR).

### Imaging

To verify the expression of GCaMP6m and the placement of the lens, mounted sections were imaged with a Zeiss LSM700 upright confocal microscope controlled by the ZEN Black acquisition software (Zeiss). Images were acquired with a 20x (PLAN APO, 0.8NA, air) or 40x (PLAN APO, 1.3NA, oil) objectives. Fixed wavelength (405nm, 488nm) lasers were used to visualize DAPI and GCaMP6m, respectively. Simultaneous differential interference contrast imaging was used to visualize the anatomical landmarks. Images were processed with FIJI.

### Analysis

*Freezing*: The proportion of time mice spent freezing in the baseline contextual freezing window as well as during tone presentations were analyzed with a custom-written script in MATLAB (Mathworks) and verified visually with the behavioral video.

*Calcium Imaging:* Acquired calcium recordings were first processed with the Inscopix Data Processing Software (IDPS) to extract calcium transients from unique individual cells. The video is pre-processed, spatially filtered, and motion corrected. Individual traces were extracted from the video with the principal component (PCA) and independent component (ICA) analysis algorithms with no downsampling. Traces with abnormal physiological transients (ie. peaks lasting over minutes) or abnormal maximal intensity projection visualizations (ie. multiple cells) were excluded. For the identification of longitudinally tracked cells, processed traces and cell maps across all analyzed days (pre-conditioning P, conditioning day 1 C1, conditioning day 5 C5, and test day T) were longitudinally registered with a spatial alignment algorithm and manually verified.

Files containing the individual traces and the general-purpose input/output (GPIO) traces were extracted and fed through a custom-written script in MATLAB. Calcium transients were denoised, normalized (per trace per day), and then binned (500ms). Tone (CS− and CS+) presentations were identified through timestamps recorded in the corresponding GPIO traces and the normalized activity in the surrounding windows (10s before tone and 45s after) were extracted. Within day average traces were generated for each cell and tone. To characterize if a cell is responsive to a tone, the average activity during the baseline (10s before onset) window is compared (Wilcoxon’s rank sum test) with that of the tone (only the first 10s is considered for pre-conditioning day). Cells are classified as either excited (p< 0.05, tone baseline delta > 0), inhibited (p< 0.05, tone baseline delta < 0), or non-responsive (p> 0.05). Shock responses were characterized in the manner by comparing activity during the baseline (4s before CS+ onset) window and a 4s period after shock onset (shock analysis window). To take into account a minority of transient shock responses where the majority of rise as well as decay were captured within the analysis window, a shortened baseline window (2s) was compared with the same period within the shock analysis window in a rolling manner. To determine if cells can differentiate CS+ from CS−, comparisons were made between the two tone windows.

In addition, to identify whether the observed responses could be discriminated from spontaneous fluctuations intrinsic to the recorded cell, the distributions of activity within all cue and baseline (or another cue) windows were concatenated and analyzed using the receiver operating characteristic analysis to calculate the discrimination index (DI = (Area Under the ROC curve – 0.5) x 2) between the cue and the baseline. To model spontaneous fluctuations, the calcium trace was circularly shifted 1000 times by a random integer and DIs were similarly calculated for each cue. Cue responses that can be positively (DI > 97.5^th^ percentile of shifted DIs), negatively (DI < 2.5^th^ percentile of shifted DIs), or not discriminated from spontaneous fluctuations were identified (α = 0.05).

To investigate whether the CS+ responses throughout conditioning can become shifted towards that of the US (FS), the similarity between each CS+ and US pairing within each conditioning day is compared by examining the Euclidean distance (ED) between the two (shock analysis window and the first 4s of CS+ window). To examine whether this shift could be consolidated across days, the day average ED were compared across days. In order to verify that potential changes in ED across pairings or across days are not influenced by random fluctuation, randomly shifted (1000 times) ED were calculated for each pairing and comparisons were made between the data-driven and random ED within and across conditioning days.

For populational analyses, cells were pooled across mice within days and analyzed. Analyzed data are stratified between mice that developed high (FG, >50% freezing to CS− on T) and low (FnG, <50% freezing to CS− on T) levels of fear generalization. For longitudinally tracked cells, cells were first characterized by their response to cues on the last day they were presented (T for tones and C5 for FS) and traced back in time.

### Statistics

All datasets were tested for normality using the Shapiro-Wilk test and found to be not normally distributed. Therefore, non-parametric comparisons were used. Statistical comparisons for the contextual freezing across days in all mice were conducted with Friedman test with post-hoc Dunn’s multiple comparison tests. Comparisons between contextual freezing across FG and FnG mice were made with Mann-Whitney tests on each day individually. Tones evoked freezing (and delta CS+ and CS− freezing) across days were analyzed with Friedman test with post-hoc Dunn’s multiple comparison tests and compared between each other using Wilcoxon signed rank test. Comparisons between delta CS+ to CS− freezing between FG and FnG mice were made with the Mann-Whitney test. Comparisons of tone (both CS− and CS+) response amplitude and DI across days in the populational analyses were made with the Kruskal-Wallis test for each type of response individually and with Friedman test for the longitudinally tracked cells. Similar comparisons for shock responses were conducted with unpaired Mann-Whitney and Wilcoxon signed rank tests. All comparisons involving the Euclidean distance between CS+ and FS were made in the same manner as tone responses. All tests were conducted with two-tails unless otherwise noted. The significance level is always set at 0.05. All statistical comparisons (except the classification of cell responses using the Wilcoxon Ranksum test, computed with MATLAB) were performed by Prism (Graphpad).

## Acknowledgements

We thank the past and current members of the Tan Lab for constructive input, the Biozentrum Imaging Facility for microscopy training, and the Inscopix team for technical support in regards to the calcium imaging experiments.

## Author Contributions

Z. Li, G. Rizzi, and K.R. Tan all contributed to the experimental design. Z. Li and G.Rizzi conducted the calcium imaging experiments. Z. Li performed all analyses and prepared the manuscript. K.R. Tan provided supervision and corrected the manuscript.

## Materials & Correspondence

All requests for materials and code should be addressed to K.R. Tan.

## Figure Legends

**Figure S1.**
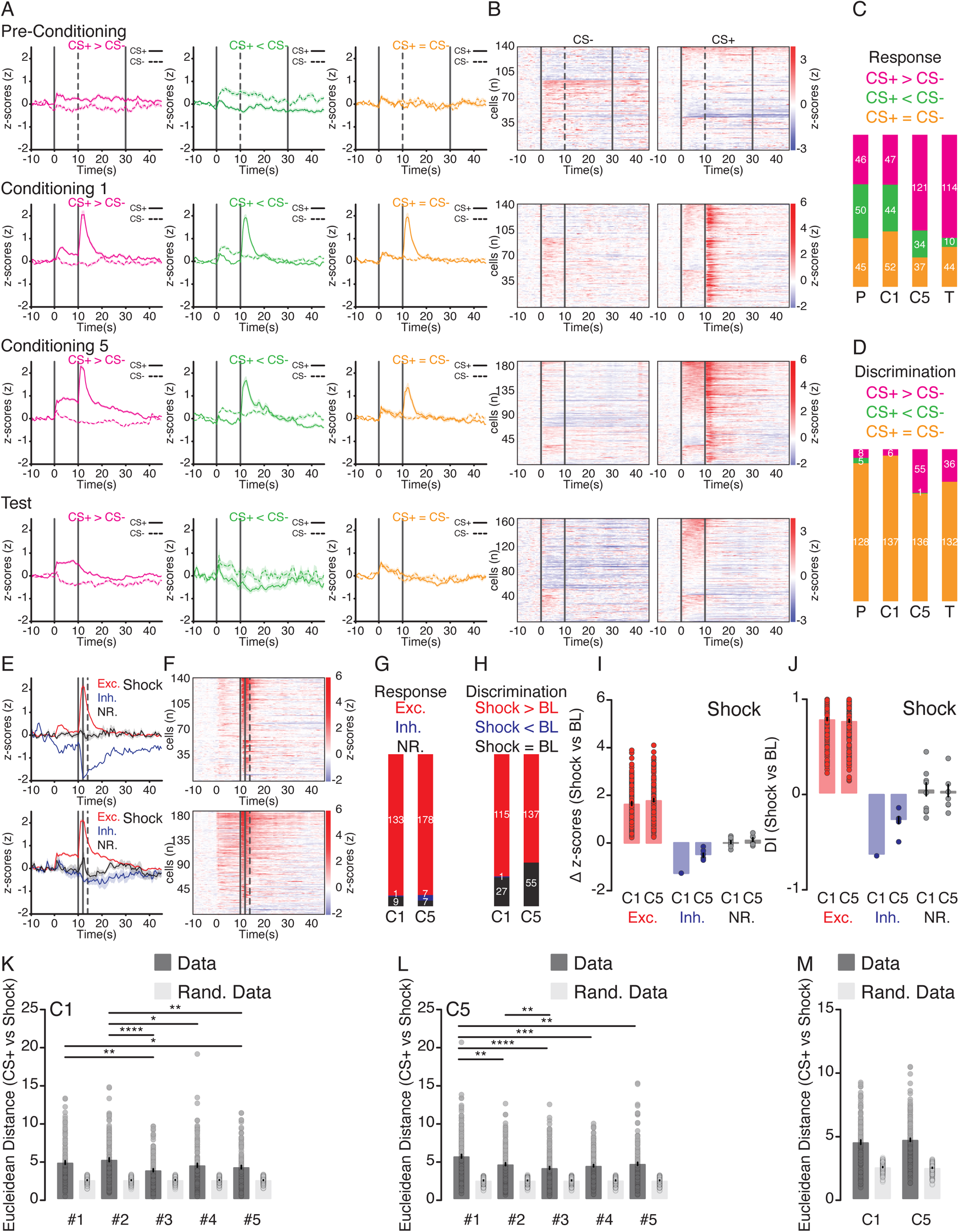
On the population level, the representations of CS+ and CS− become more distinguishable while that of the CS+ and footshocks become more similar. A) Average traces of cells that are differentially modulated by CS+ and CS− in P, C1, C5, and T days (black bars denote tone windows). B) Heatmaps of cell responses to CS+ and CS−, sorted by the difference between CS+ and CS− modulations in P, C1, C5, and T days. C) Stacked bar chart of proportion of cells differentially modulated by CS+ as compared to CS− in P, C1, C5, and T days. D) Stacked bar chart of proportion of cells whose discrimination between CS+ and CS− is significantly distinct from intrinsic activity fluctuations in P, C1, C5, and T days. E) Average traces of shock responsive cells in C1 and C5 days (solid and dotted bar denotes shock and shock analysis windows, respectively). F) Heatmaps of shock responses in C1 and C5. G) Stacked bar chart showing proportion of shock responses in C1 and C5. H) Stacked bar chart of shock responses that can be discriminated from spontaneous activity in C1 and C5. I) Bar plot depicting the amplitude of shock responses in C1 and C5. J) Bar plot denoting the DI of shock responses in C1 and C5. K) Bar graph with the Euclidean distance between CS+ and FS pairings within C1. L) Bar graph of the Euclidean distance between CS+ and FS pairings within C5. M) Bar graph presenting the average Euclidean distance between CS+ and FS pairings in C1 and C5. Data are presented as mean ± SEM *p<0.05, **p<0.01, ***p<0.001, ****p<0.0001, 81-141-192 cells from 8 mice.

**Figure S2.**
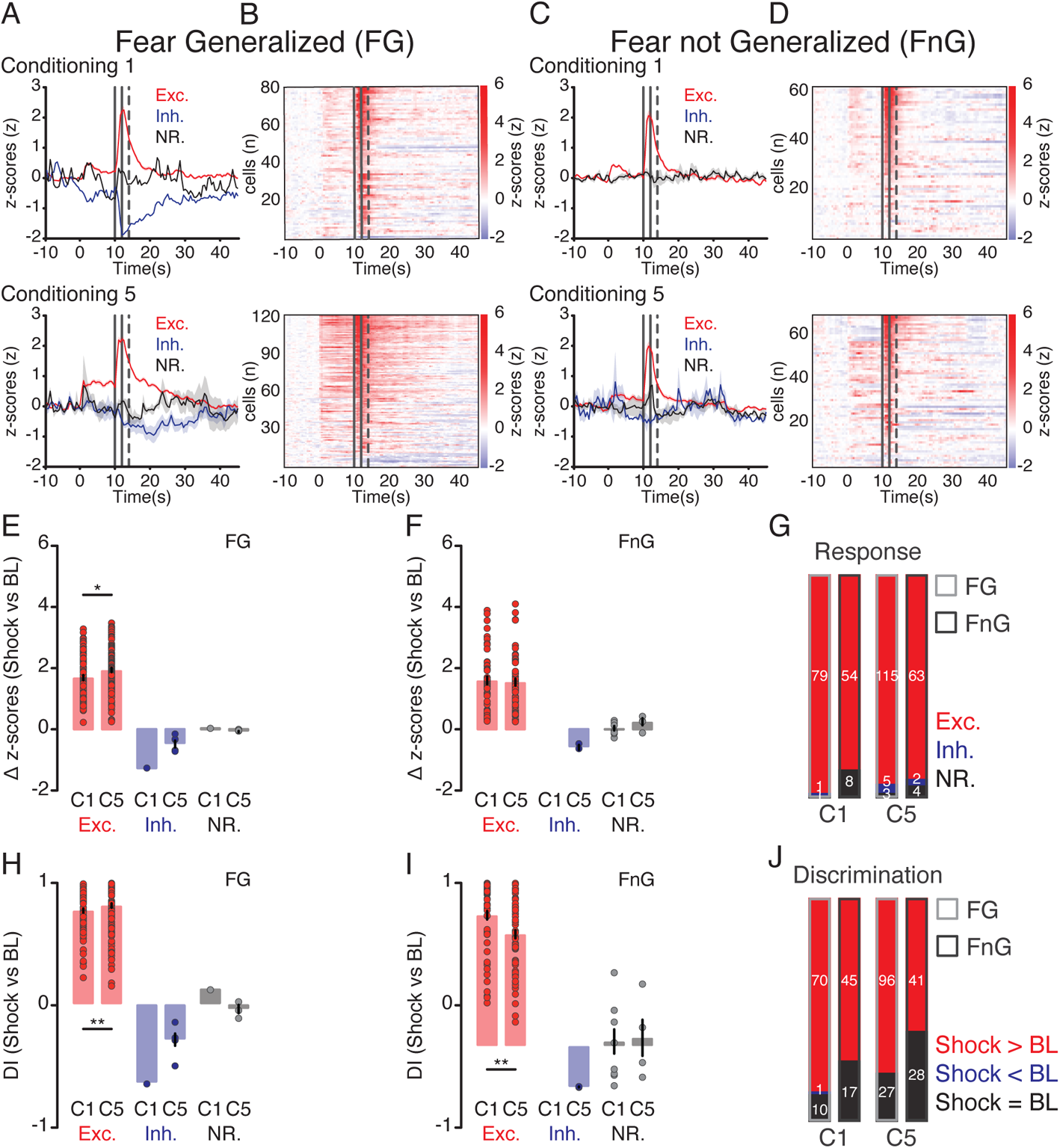
ZI neurons of both fear generalized and not generalized mice are strongly excited by footshocks. A) Average traces of shock responses (solid and dotted lines denote shock and shock analysis windows) of fear generalized mice in C1 and C5 days. B) Heatmaps of shock responses from fear generalized mice in C1 and C5 days. C) Average traces of shock responses in mice that did not generalized fear on C1 and C5. D) Heatmaps of shock responses in mice that did not generalize fear on C1 and C5. E) Bar graph summarizing the amplitude of shock responses from fear generalized mice on C1 and C5 days. F) Bar graph of the amplitude of shock responses from mice that did not generalized fear on C1 and C5. G) Stacked bar chart showing the proportion of shock responsive cells between fear generalized and not generalized mice on C1 and C5. H) Bar graph with the DI of shock responses in fear generalized mice on C1 and C5 days. I) Bar graph depicting the DI of shock responses in mice that did not generalized fear on C1 and C5. J) Stacked bar chart highlighting the proportion of shock responses that can be discriminated from spontaneous activity in mice that generalized and did not generalized fear on C1 and C5. Data are presented as mean ± SEM *p<0.05, **p<0.01, ***p<0.001, ****p<0.0001, 81-123 cells (fear generalized, 3 mice) and 48-69 cells (fear not generalized, 5 mice).

**Figure S3.**
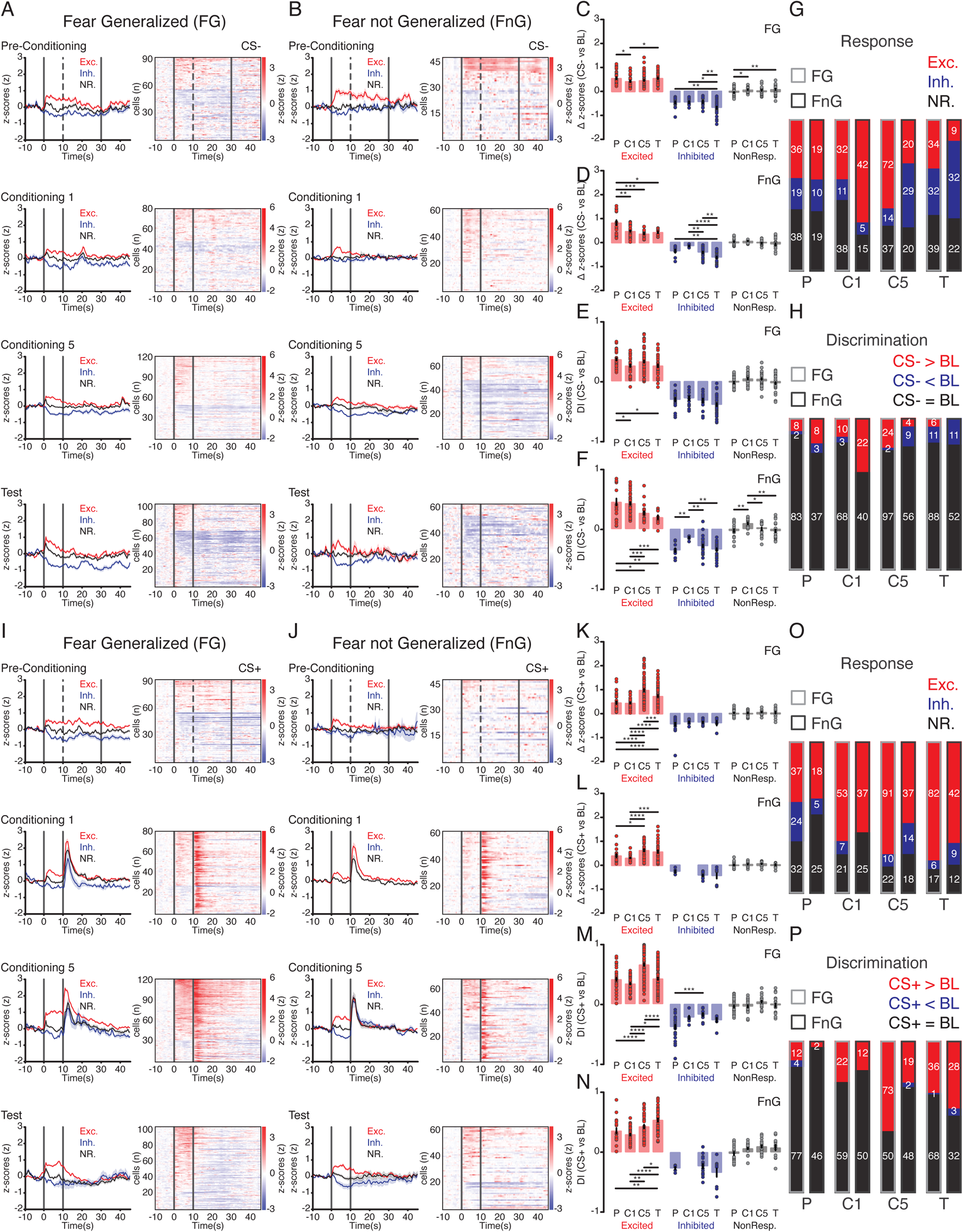
ZI neurons of fear generalized and not generalized mice encode CS+ similarly but differentially encode CS−. A) Average traces and heatmaps of CS+ responses from fear generalized mice in P, C1, C5, T days (black bars depict tone windows). B) Average traces and heatmaps of CS+ responses from mice that did not generalized fear. C) Bar graph with the amplitude of CS+ responses from mice that generalized fear. D) Bar graph with the DI of CS+ responses of mice that generalized fear. E) Bar graph with the amplitude of CS+ responses from mice that did not generalized fear. F) Bar graph with the DI of CS+ responses of mice that did not generalized fear. G) Stacked bar chart with proportion of CS+ responsive cells between fear generalized and not generalized mice. H) Stacked bar chart with proportion of cells that can discriminate CS+ responses from spontaneous activity fluctuations between fear generalized and not generalized mice. I) Average traces and heatmaps of CS− responses from fear generalized mice in P, C1, C5, T days. J) Average traces and heatmaps of CS− responses from mice that did not generalized fear. K) Bar graph with the amplitude of CS− responses from mice that generalized fear. L) Bar graph with the DI of CS− responses of mice that generalized fear. M) Bar graph with the amplitude of CS− responses from mice that did not generalized fear. N) Bar graph with the DI of CS− responses of mice that did not generalized fear. O) Stacked bar chart with proportion of CS− responsive cells between fear generalized and not generalized mice. P) Stacked bar chart with proportion of cells that can discriminate CS− responses from spontaneous activity fluctuations between fear generalized and not generalized mice. Data are presented as mean ± SEM *p<0.05, **p<0.01, ***p<0.001, ****p<0.0001, 81-123 cells (fear generalized, 3 mice) and 48-69 cells (fear not generalized, 5 mice).

**Figure S4.**
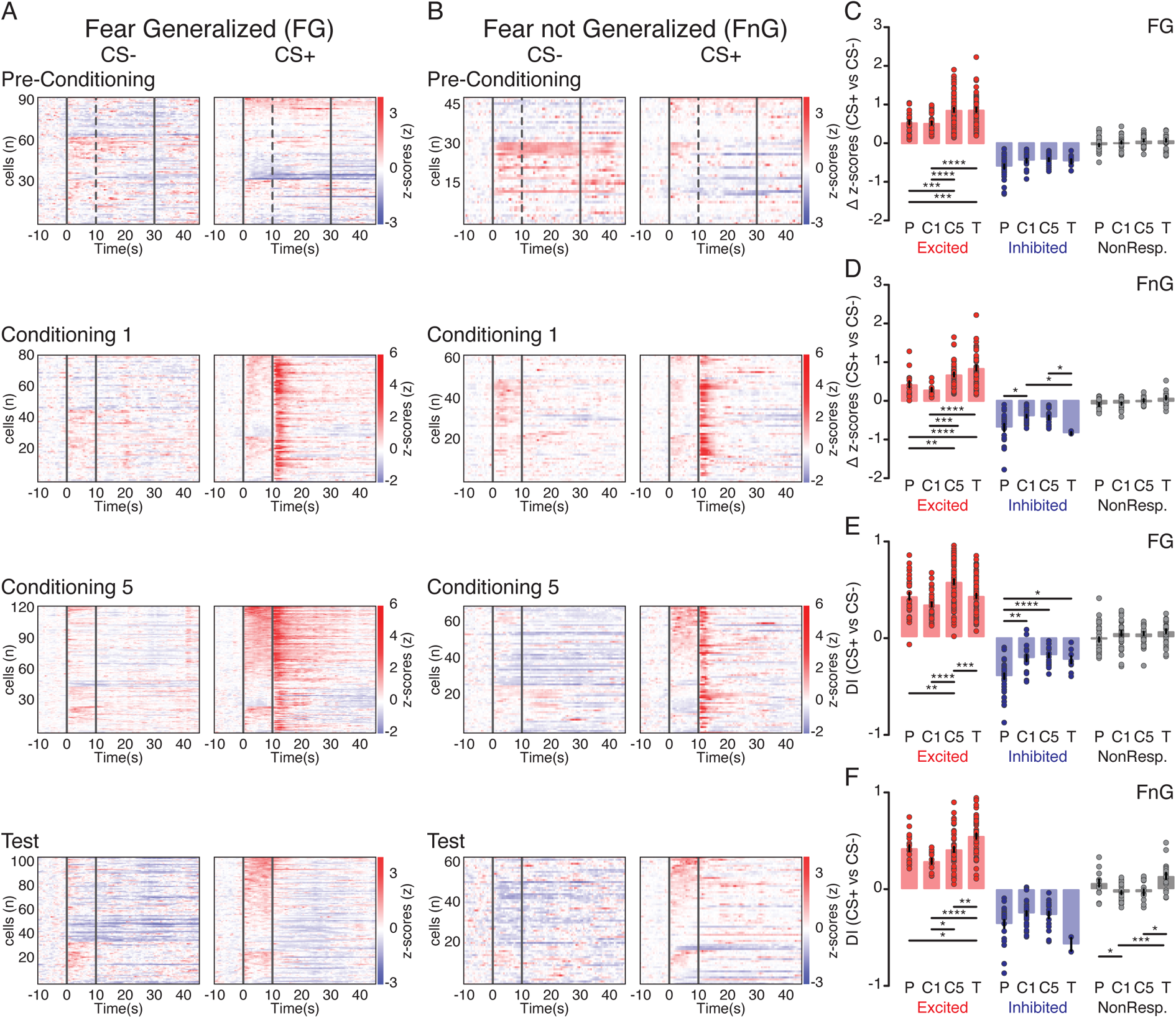
The magnitude of positive discrimination between CS+ and CS− is potentiated in the ZI neurons of both fear generalized and not generalized mice. A) Heatmaps of CS+ and CS− responses from fear generalized mice in P, C1, C5, and T days, sorted by magnitude of differences between CS+ and CS− modulations (black bars highlight tone windows). B) Heatmaps of CS+ and CS− responses from mice that did not develop fear generalization in P, C1, C5, and T days, sorted by magnitude of differences between CS+ and CS− modulations. C) Bar graph of differences in CS+ and CS− modulations of fear generalized mice in P, C1, C5, and T days. D) Bar graph of differences in activity between CS+ and CS− observed from mice that did not develop fear generalization in P, C1, C5, and T days. E) Bar graph of DI between CS+ and CS− responses in fear generalized mice on P, C1, C5, and T days. F) Bar graph of DI between CS+ and CS− responses in mice that did not generalized fear on P, C1, C5, and T days. Data are presented as mean ± SEM *p<0.05, **p<0.01, ***p<0.001, ****p<0.0001, 81-123 cells (fear generalized, 3 mice) and 48-69 cells (fear not generalized, 5 mice).

**Video S1. Example behavioral recording of a mice performing in the fear conditioning task on the C1 with the first CS− and CS+ presentations (2x speed).**

**Video S2. Example behavioral recording of a mice performing in the fear conditioning task on the T responding to the CS− (2x speed).**

**Video S3. Example behavioral recording of a mice performing in the fear conditioning task on the T responding to the CS+ (2x speed).**

**Video S4. Example calcium imaging recording of ZI neurons while the recorded mouse was presented with CS−, and CS+ and FS on the C5 (2x speed).**

**Video S5. Example simultaneous behavioral recording of a FG mouse performing in the fear conditioning task on the T responding to the CS− and CS+ with the concomitant calcium imaging recording below (2x speed).**

**Video S6. Example simultaneous behavioral recording of a FnG mouse performing in the fear conditioning task on the T responding to the CS− and CS+ with the concomitant calcium imaging recording below (2x speed).**

**Table S1.**
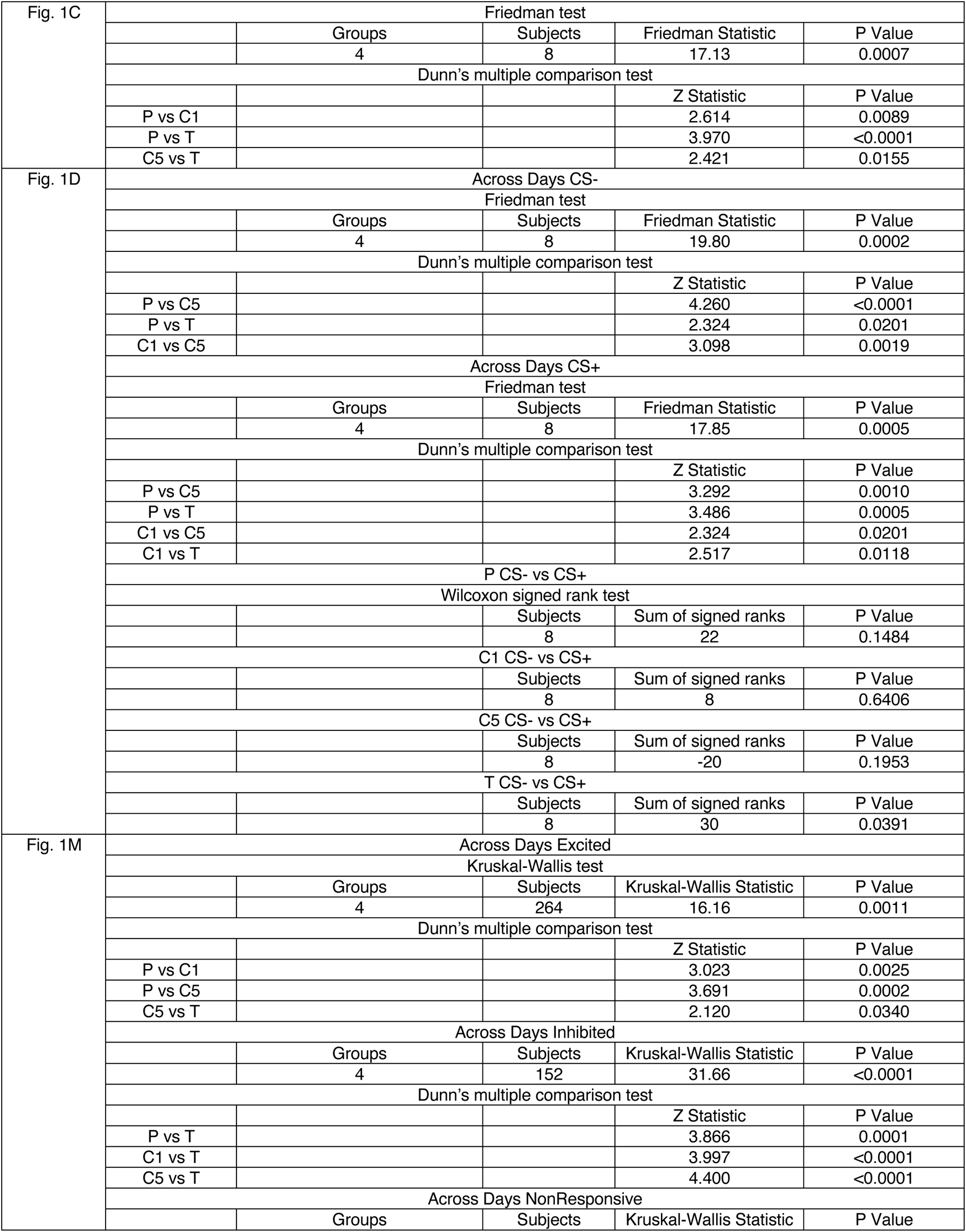
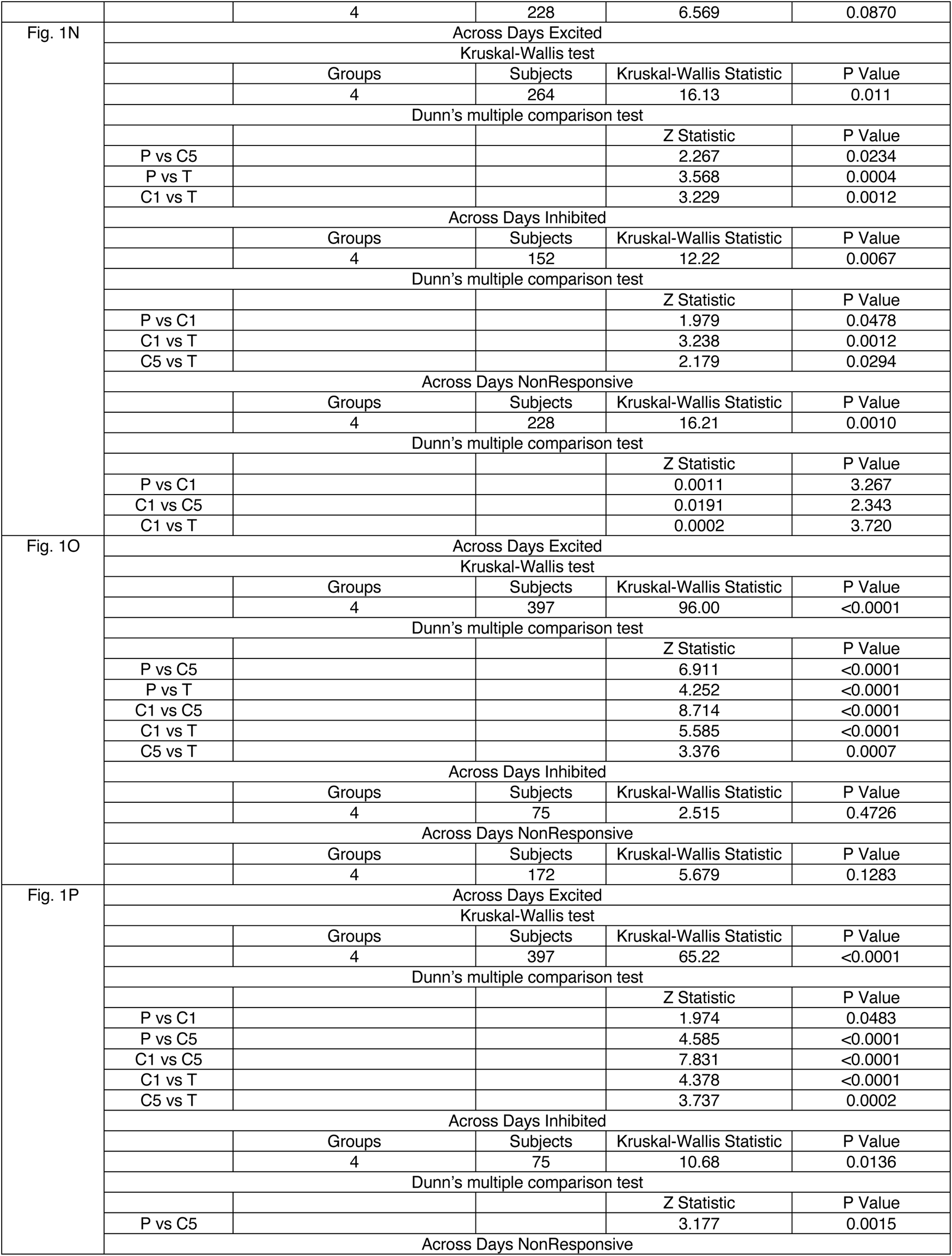
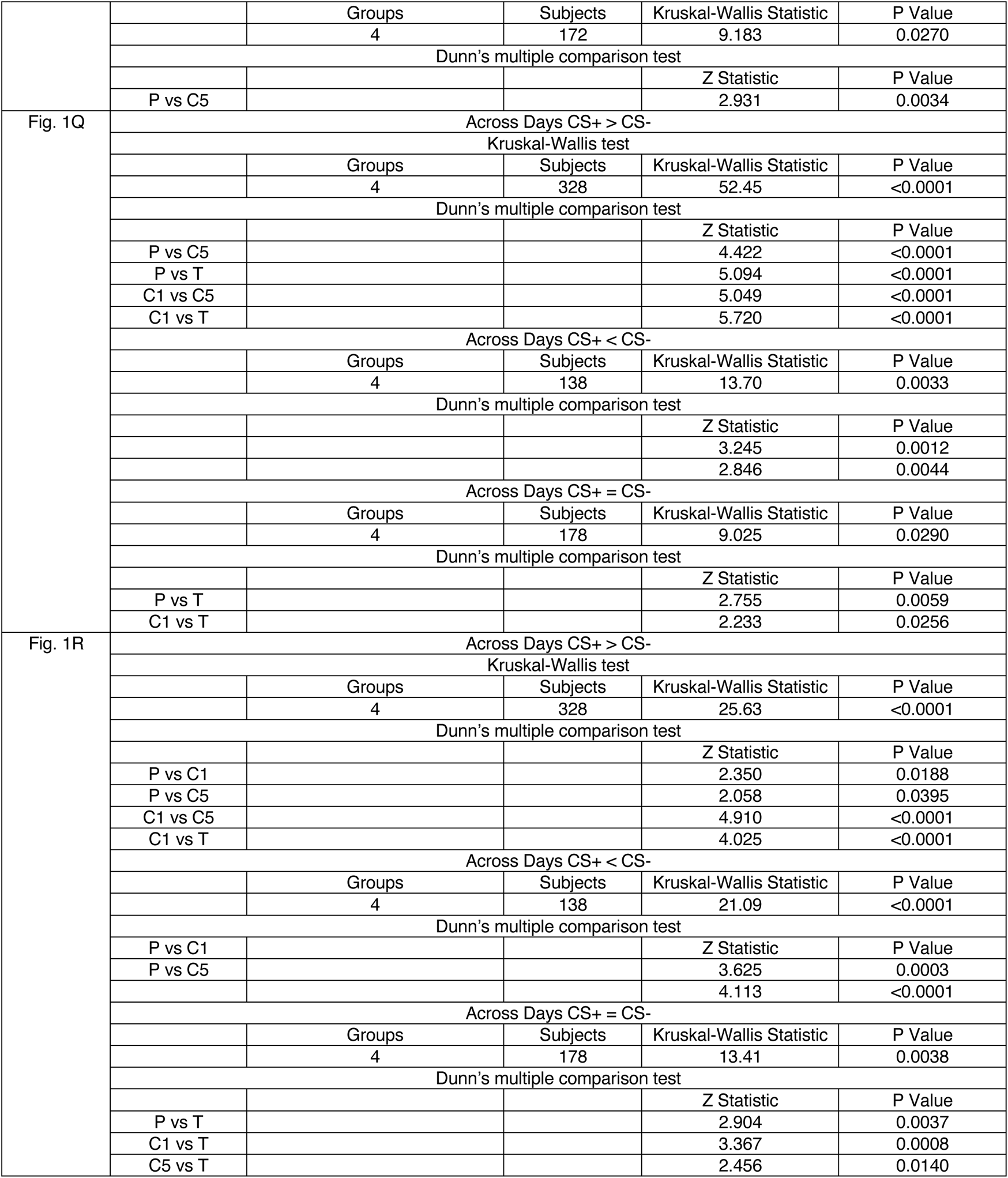
Statistical Information for Figure 1

**Table S2.**
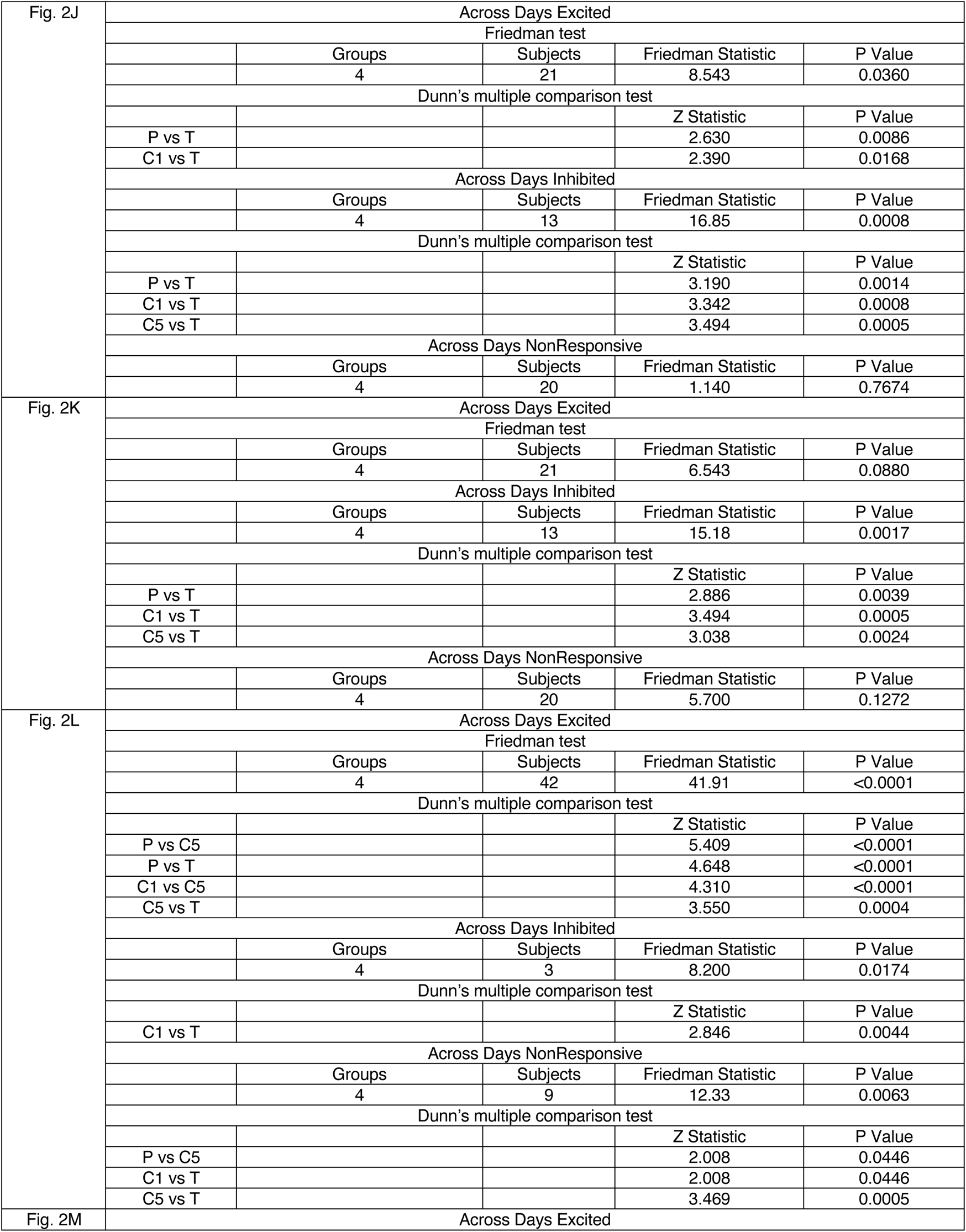
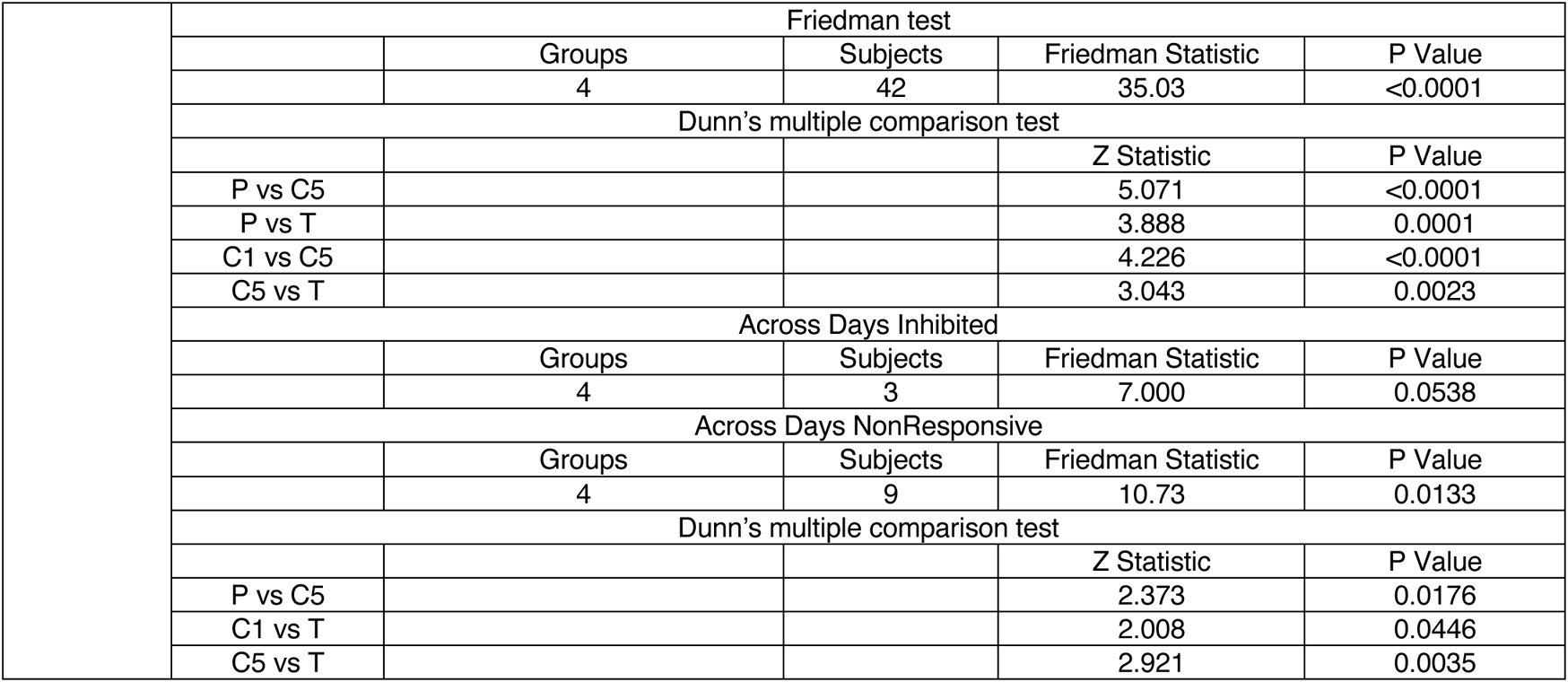
Statistical Information for Figure 2

**Table S3.**
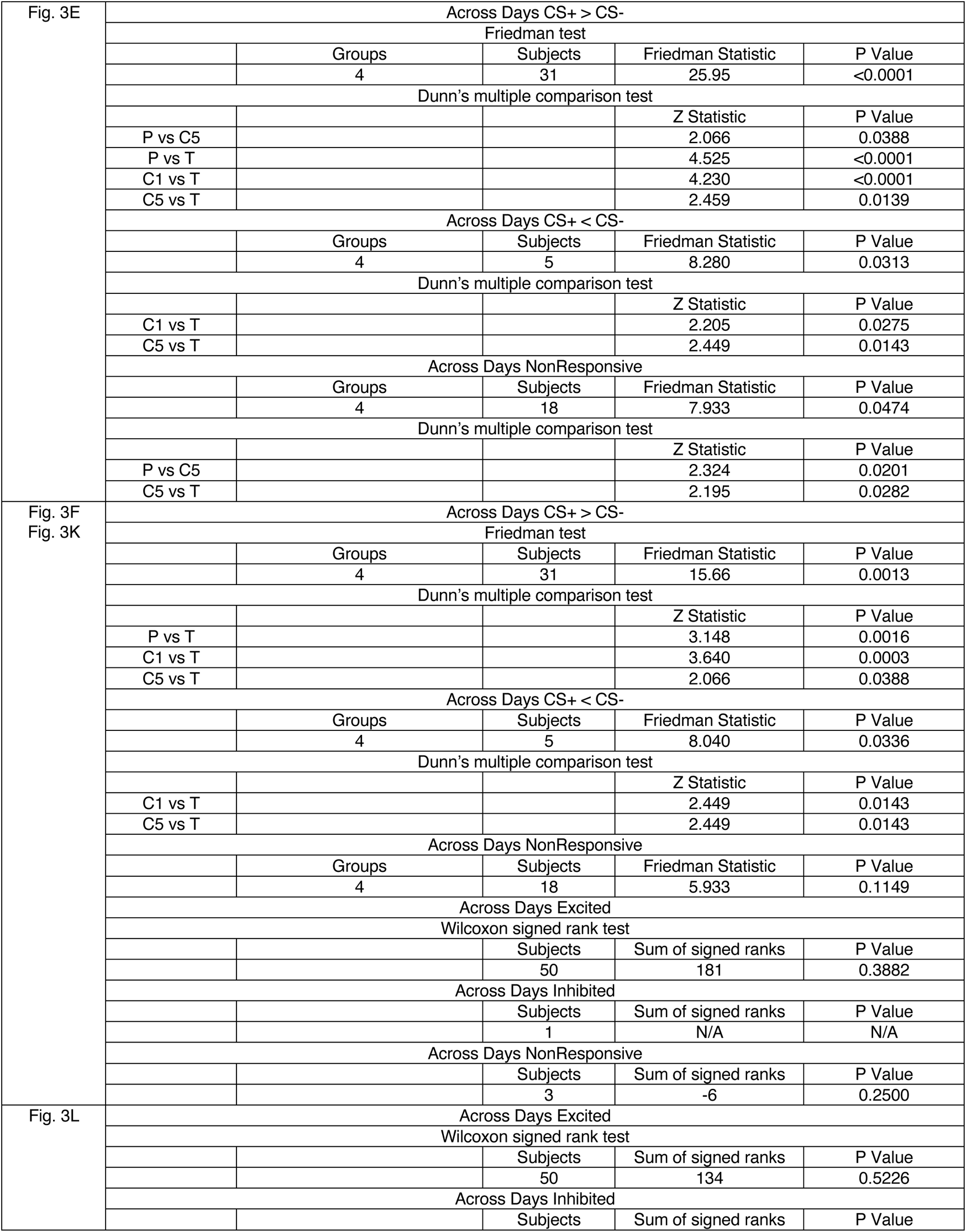
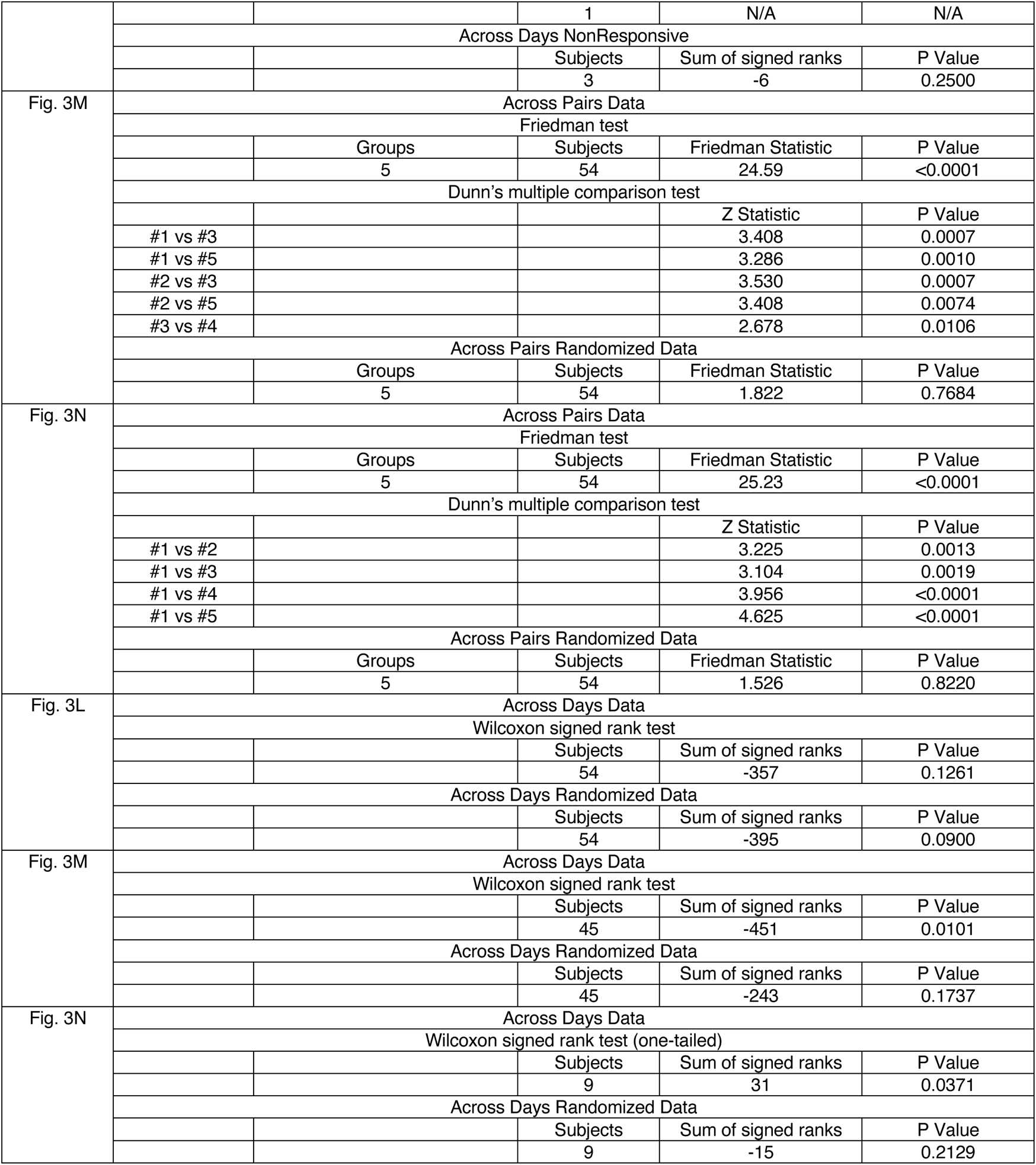
Statistical Information for Figure 3

**Table S4.**
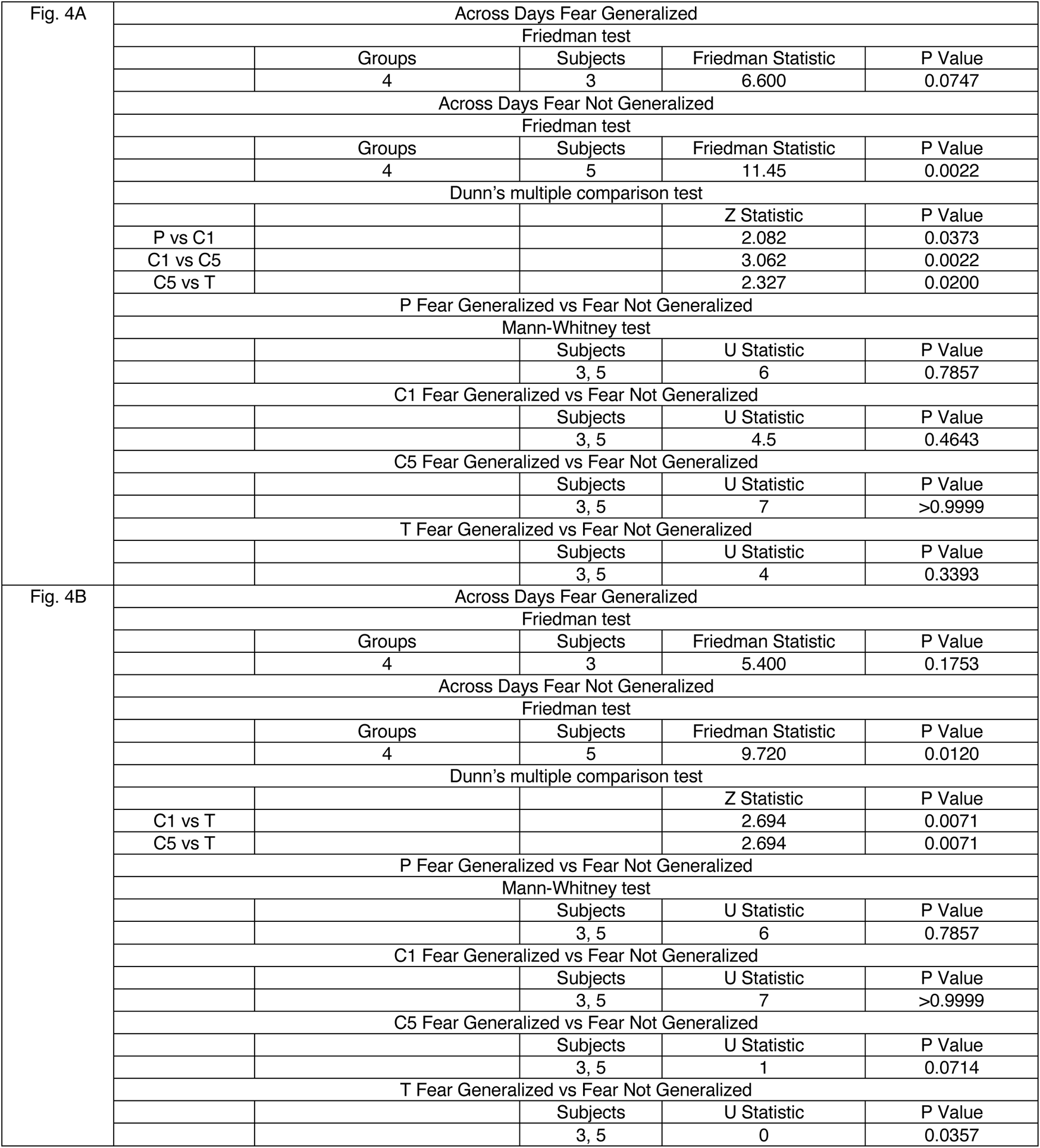
Statistical Information for Figure 4

**Table S5.** Statistical Information for Figure S1

|  |  |  |  |  |
| --- | --- | --- | --- | --- |
| Fig. S1I | Across Days Excited |  |  |  |
|  | Mann-Whitney test |  |  |  |
|  |  | Subjects | U Statistic | P Value |
|  |  | 133, 178 | 10562 | 0.1043 |
|  | Across Days Inhibited |  |  |  |
|  |  | Subjects | U Statistic | P Value |
|  |  | 1,7 | N/A | N/A |
|  | Across Days NonResponsive |  |  |  |
|  |  | Subjects | U Statistic | P Value |
|  |  | 9,7 | 21 | 0.2991 |
| Fig. S1J | Across Days Excited |  |  |  |
|  | Mann-Whitney test |  |  |  |
|  |  | Subjects | U Statistic | P Value |
|  |  | 133, 178 | 11738 | 0.8986 |
|  | Across Days Inhibited |  |  |  |
|  |  | Subjects | U Statistic | P Value |
|  |  | 1,7 | N/A | N/A |
|  | Across Days NonResponsive |  |  |  |
|  |  | Subjects | U Statistic | P Value |
|  |  | 9,7 | 31 | >0.9999 |
| Fig. S1K | Across Pairs Data |  |  |  |
|  | Kruskal-Wallis test |  |  |  |
|  | Groups | Subjects | Friedman Statistic | P Value |
|  | 5 | 710 | 19.65 | 0.0006 |
|  | Dunn's multiple comparison test |  |  |  |
|  |  |  | Z Statistic | P Value |
|  | #1 vs #3 |  | 3.086 | 0.0020 |
|  | #1 vs #5 |  | 2.014 | 0.0440 |
|  | #2 vs #3 |  | 3.919 | <0.0001 |
|  | #2 vs #4 |  | 2.246 | 0.0247 |
|  | #2 vs #5 |  | 2.841 | 0.0045 |
|  | Across Pairs Randomized Data |  |  |  |
|  | Groups | Subjects | Friedman Statistic | P Value |
|  | 5 | 710 | 0.1559 | 0.9971 |
| Fig. S1L | Across Pairs Data |  |  |  |
|  | Kruskal-Wallis test |  |  |  |
|  | Groups | Subjects | Friedman Statistic | P Value |
|  | 5 | 958 | 27.62 | <0.0001 |
|  | Dunn's multiple comparison test |  |  |  |
|  |  |  | Z Statistic | P Value |
|  | #1 vs #2 |  | 3.034 | 0.0024 |
|  | #1 vs #3 |  | 5.130 | <0.0001 |
|  | #1 vs #4 |  | 3.519 | 0.0004 |
|  | #1 vs #5 |  | 3.091 | 0.0020 |
|  | #2 vs #3 |  | 2.096 | 0.0361 |
|  | #3 vs #5 |  | 2.039 | 0.0414 |
|  | Across Pairs Randomized Data |  |  |  |
|  | Groups | Subjects | Friedman Statistic | P Value |
|  | 5 | 958 | 0.1506 | 0.9973 |
| Fig. S1M | Across Days Data |  |  |  |
|  | Mann-Whitney test |  |  |  |
|  |  | Subjects | U Statistic | P Value |
|  |  | 143, 192 | 12783 | 0.2814 |
|  | Across Days Randomized Data |  |  |  |
|  |  | Subjects | U Statistic | P Value |
|  |  | 143, 192 | 13008 | 0.4119 |

**Table S6.**
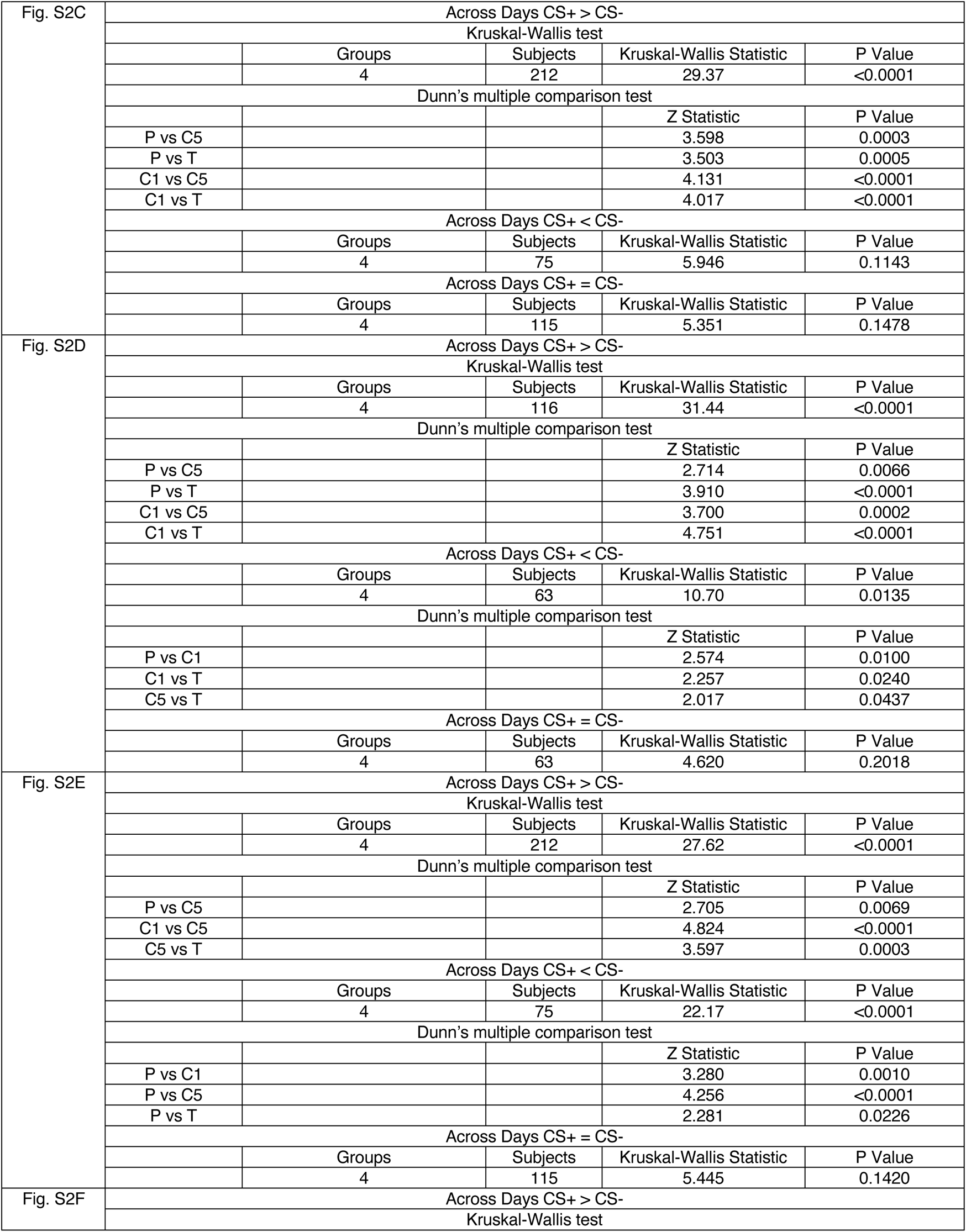
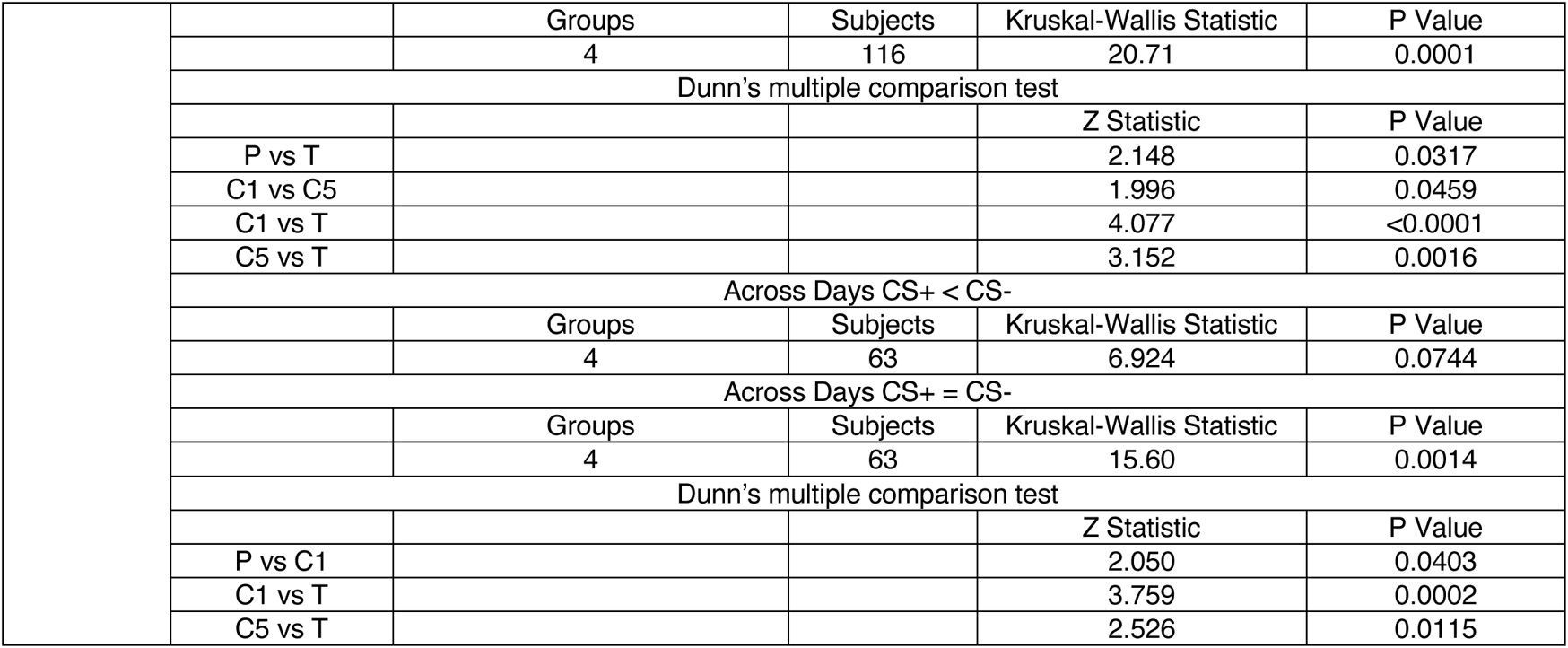
Statistical Information for Figure S2

**Table S7.**
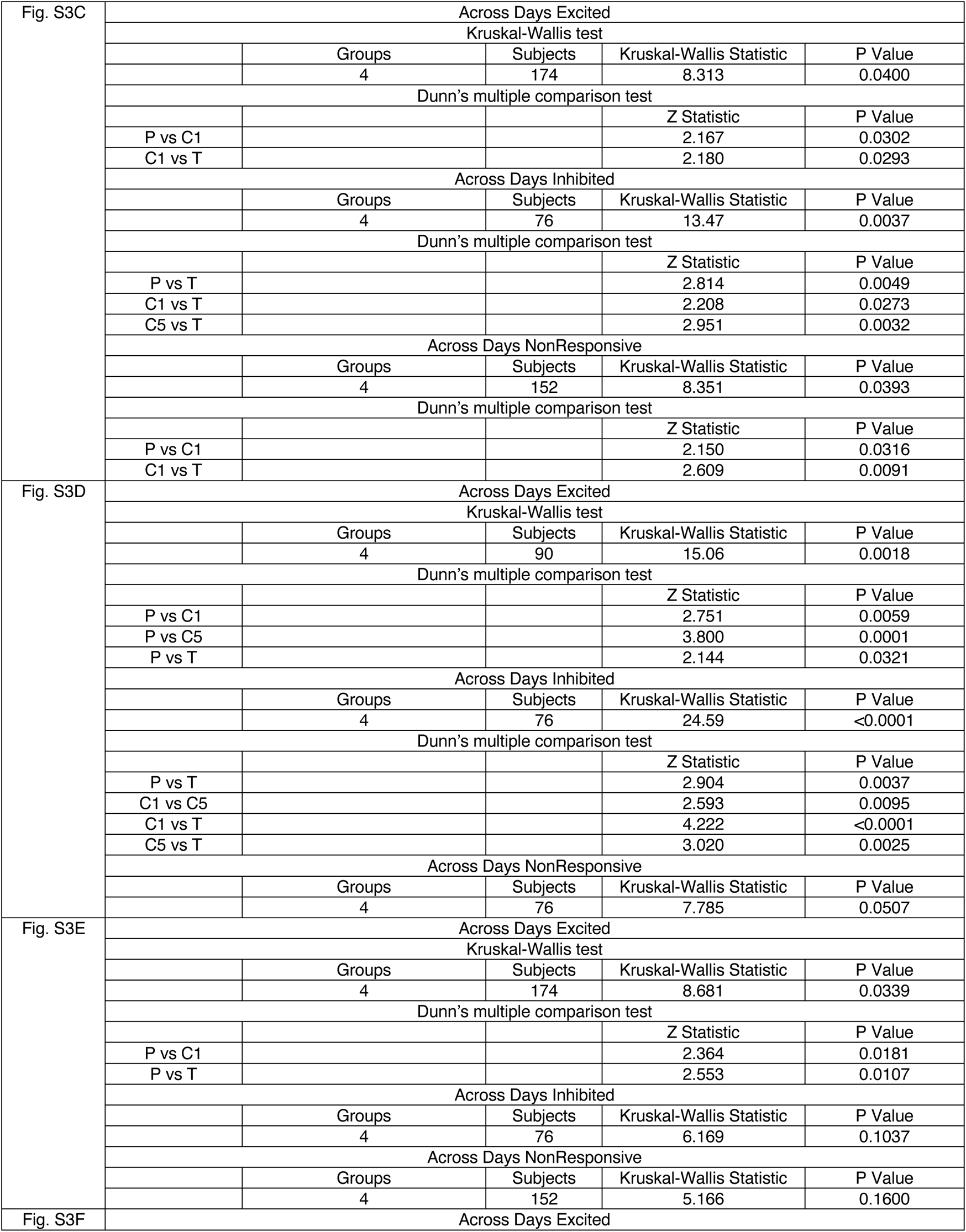
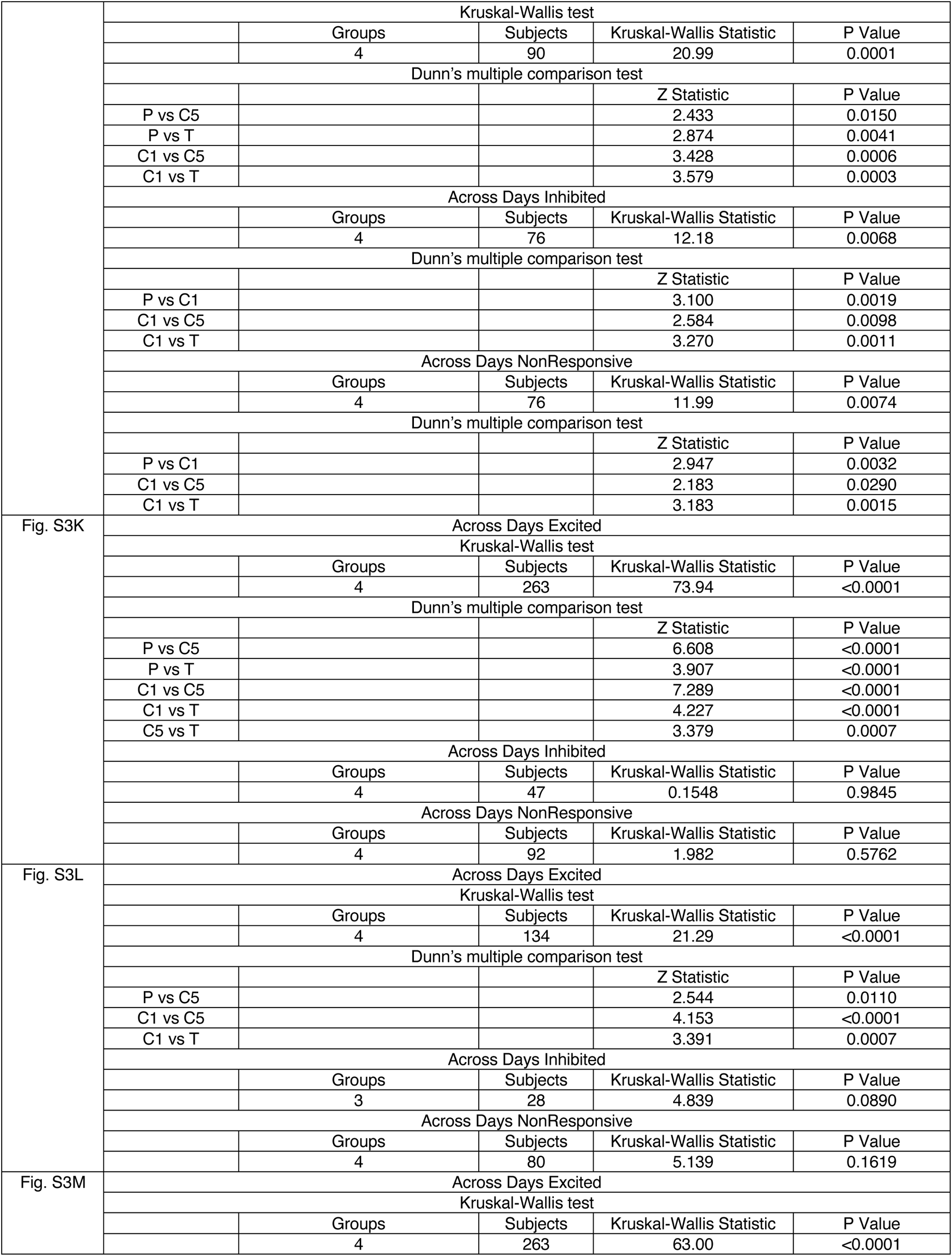
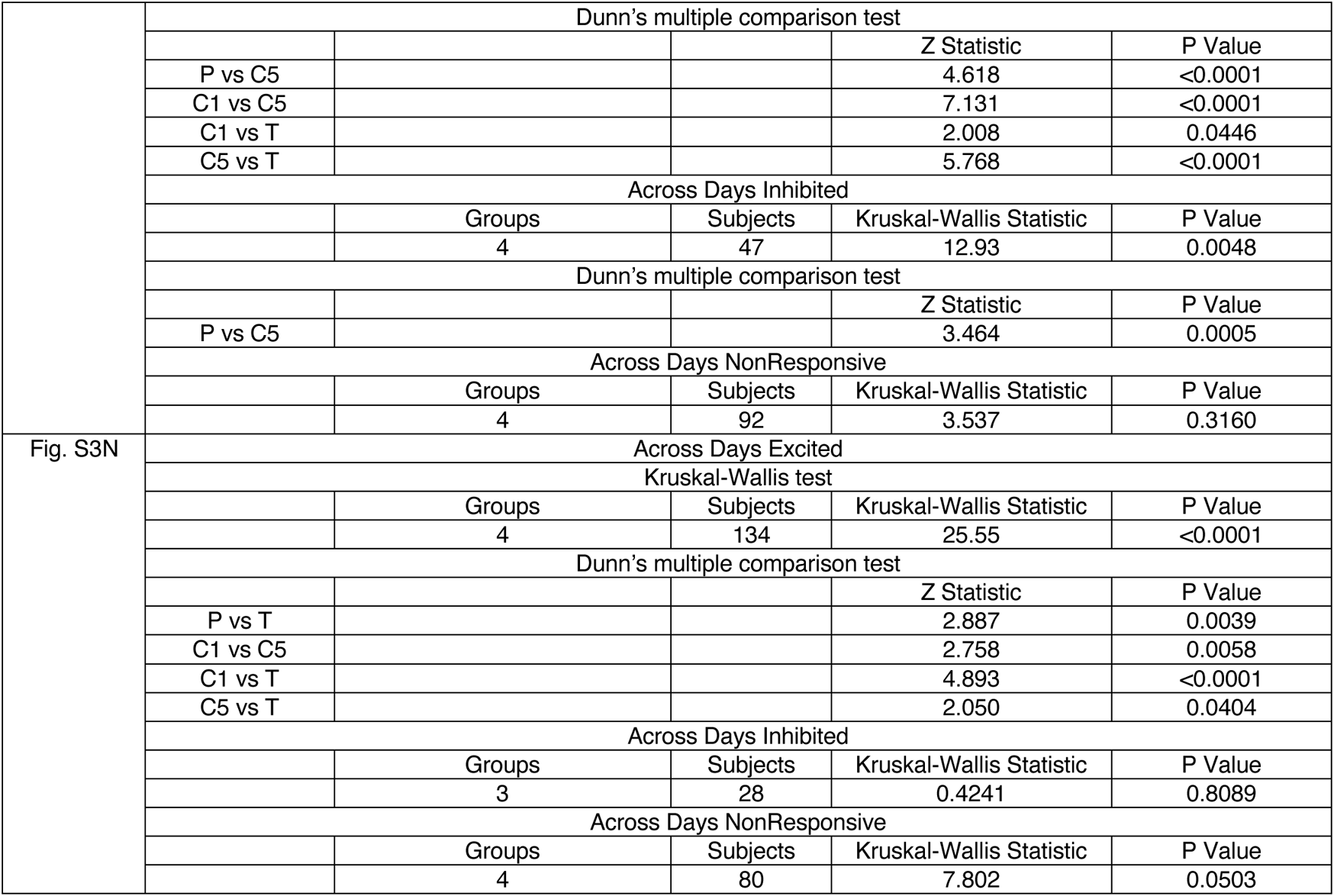
Statistical Information for Figure S3

**Table S8.** Statistical Information for Figure S4

|  |  |  |  |  |
| --- | --- | --- | --- | --- |
| Fig. S4E | Across Days Excited |  |  |  |
|  | Mann-Whitney test |  |  |  |
|  |  | Subjects | U Statistic | P Value |
|  |  | 79, 115 | 3669 | 0.0228 |
|  | Across Days Inhibited |  |  |  |
|  |  | Subjects | U Statistic | P Value |
|  |  | 1, 5 | N/A | N/A |
|  | Across Days NonResponsive |  |  |  |
|  |  | Subjects | U Statistic | P Value |
|  |  | 1, 3 | N/A | N/A |
| Fig. S4F | Across Days Excited |  |  |  |
|  | Mann-Whitney test |  |  |  |
|  |  | Subjects | U Statistic | P Value |
|  |  | 54, 63 | 1662 | 0.8339 |
|  | Across Days Inhibited |  |  |  |
|  |  | Subjects | U Statistic | P Value |
|  |  | 0, 2 | N/A | N/A |
|  | Across Days NonResponsive |  |  |  |
|  |  | Subjects | U Statistic | P Value |
|  |  | 8, 4 | 8 | 0.2141 |
| Fig. S4H | Across Days Excited |  |  |  |
|  | Mann-Whitney test |  |  |  |
|  |  | Subjects | U Statistic | P Value |
|  |  | 79, 115 | 3519 | 0.0075 |
|  | Across Days Inhibited |  |  |  |
|  |  | Subjects | U Statistic | P Value |
|  |  | 1, 5 | N/A | N/A |
|  | Across Days NonResponsive |  |  |  |
|  |  | Subjects | U Statistic | P Value |
|  |  | 1, 3 | N/A | N/A |
| Fig. S4I | Across Days Excited |  |  |  |
|  | Mann-Whitney test |  |  |  |
|  |  | Subjects | U Statistic | P Value |
|  |  | 54, 63 | 1151 | 0.0024 |
|  | Across Days Inhibited |  |  |  |
|  |  | Subjects | U Statistic | P Value |
|  |  | 0, 2 | N/A | N/A |
|  | Across Days NonResponsive |  |  |  |
|  |  | Subjects | U Statistic | P Value |
|  |  | 8, 4 | 15 | 0.9333 |

## References

1. Blanchard, D. C. & Blanchard, R. J. Chapter 2.4 Defensive behaviors, fear, and anxiety. in Handbook of Behavioral Neuroscience (eds. Blanchard, R. J., Blanchard, D. C., Griebel, G. & Nutt, D.) vol. 17 63–79 (Elsevier, 2008).

2. LeDoux, J. Rethinking the emotional brain. Neuron 73, 653–676 (2012).

3. Tovote, P., Fadok, J. P. & Lüthi, A. Neuronal circuits for fear and anxiety. Nature Reviews Neuroscience 16, 317–331 (2015).

4. Maren, S. Neurobiology of Pavlovian fear conditioning. Annu Rev Neurosci 24, 897–931 (2001).

5. Ciocchi, S. et al. Encoding of conditioned fear in central amygdala inhibitory circuits. Nature 468, 277–282 (2010).

6. Grewe, B. F. et al. Neural ensemble dynamics underlying a long-term associative memory. Nature 543, 670–675 (2017).

7. Courtin, J. et al. Prefrontal parvalbumin interneurons shape neuronal activity to drive fear expression. Nature 505, 92–96 (2014).

8. Cummings, K. A. & Clem, R. L. Prefrontal somatostatin interneurons encode fear memory. Nat Neurosci 23, 61–74 (2020).

9. Tang, J., Wagner, S., Schachner, M., Dityatev, A. & Wotjak, C. T. Potentiation of amygdaloid and hippocampal auditory-evoked potentials in a discriminatory fear-conditioning task in mice as a function of tone pattern and context. Eur J Neurosci 18, 639–650 (2003).

10. Xu, C. et al. Distinct Hippocampal Pathways Mediate Dissociable Roles of Context in Memory Retrieval. Cell 167, 961–972.e16 (2016).

11. Rizzi, G., Li, Z., Hogrefe, N. & Tan, K. R. Lateral ventral tegmental area GABAergic and glutamatergic modulation of conditioned learning. Cell Rep 34, 108867 (2021).

12. Tovote, P. et al. Midbrain circuits for defensive behaviour. Nature 534, 206– 212 (2016).

13. Chou, X.-L. et al. Inhibitory gain modulation of defense behaviors by zona incerta. Nat Commun 9, 1151 (2018).

14. Burrows, A. M. et al. Limbic and motor function comparison of deep brain stimulation of the zona incerta and subthalamic nucleus. Neurosurgery 70, 125– 130; discussion 130-131 (2012).

15. Zhou, M. et al. A central amygdala to zona incerta projection is required for acquisition and remote recall of conditioned fear memory. Nat. Neurosci. 21, 1515–1519 (2018).

16. Asok, A., Kandel, E. R. & Rayman, J. B. The Neurobiology of Fear Generalization. Front Behav Neurosci 12, 329 (2018).

17. Venkataraman, A. et al. Modulation of fear generalization by the zona incerta. Proc. Natl. Acad. Sci. U.S.A. 116, 9072–9077 (2019).

18. Taylor, J. A. et al. Single cell plasticity and population coding stability in auditory thalamus upon associative learning. Nat Commun 12, 2438 (2021).

19. MacNamara, A. et al. Neural correlates of individual differences in fear learning. Behav Brain Res 287, 34–41 (2015).

20. Bush, D. E. A., Sotres-Bayon, F. & LeDoux, J. E. Individual differences in fear: isolating fear reactivity and fear recovery phenotypes. J Trauma Stress 20, 413– 422 (2007).

21. Graham, B. M. & Richardson, R. Individual differences in the expression of conditioned fear are associated with endogenous fibroblast growth factor 2. Learn Mem 23, 42–45 (2016).

22. Duvarci, S., Bauer, E. P. & Paré, D. The bed nucleus of the stria terminalis mediates inter-individual variations in anxiety and fear. J Neurosci 29, 10357– 10361 (2009).

23. Loskutova, L. V., Vinnitskii, I. M. & Il’yuchenok, R. Yu. Participation of the zona incerta in mechanisms of the conditioned avoidance reaction. Neurosci. Behav. Physiol. 11, 60–62 (1981).

24. Liu, K. et al. Lhx6-positive GABA-releasing neurons of the zona incerta promote sleep. Nature 548, 582–587 (2017).

25. Dymond, S., Dunsmoor, J. E., Vervliet, B., Roche, B. & Hermans, D. Fear Generalization in Humans: Systematic Review and Implications for Anxiety Disorder Research. Behavior Therapy 46, 561–582 (2015).

26. Stegmann, Y. et al. Individual differences in human fear generalization-pattern identification and implications for anxiety disorders. Transl Psychiatry 9, 307 (2019).

27. Keith, F. B. J. & Paxinos, G. The Mouse Brain in Stereotaxic Coordinates. (Elsevier Inc., 2007).

